# BCL11b interacts with RNA and proteins involved in RNA processing and developmental diseases

**DOI:** 10.1101/2020.11.05.369959

**Authors:** Haitham Sobhy, Marco De Rovere, Amina Ait-Ammar, Muhammad Kashif, Clementine Wallet, Fadoua Daouad, Thomas Loustau, Carine Van Lint, Christian Schwartz, Olivier Rohr

**Author notes:** Correspondence should be addressed to Haitham Sobhy and Olivier ROHR. HS, MDR and AAA can be considered as equal contributors. CVL, CS and OR can be considered as equal contributors.

## Abstract

BCL11b (B-cell lymphoma/leukemia 11B, CTIP2) is a C2H2 zinc-finger transcription regulator and tumor suppressor. BCL11b is involved in lymphomagenesis and in fetal, central nervous system (CNS) and immune system developments. Therefore, it may contribute in congenital disorders and cancers (e.g. leukemia). BCL11b favors persistence of HIV latency in microglia, CNS macrophages. BCL11b contributes to control cell cycle, differentiation and apoptosis in multiple organisms and cell models; however exact mechanisms are unknown. Although BCL11b recruits non-coding RNA and epigenetic enzymes to regulate gene expression, BCL11b-associated ribonucleoprotein complexes are unknown. Here, using immunoprecipitation of BCL11b-binding RNA and proteins (CLIP-seq and quantitative LC-MS/MS mass spectrometry) complemented with systems biology validations, we show that BCL11b interacts with RNA splicing and nonsense-mediated decay proteins, including FUS, SMN1, UPF1 and Drosha, which may contribute in isoform selection of protein-coding RNA isoforms from noncoding-RNAs isoforms (retained introns or nonsense mediated RNA). Interestingly, BCL11b binds to RNA transcripts and proteins encoded by the same genes (FUS, ESWR1, CHD and Tubulin). Our study highlights that BCL11b targets RNA processing and splicing proteins, and RNAs that implicate cell cycle, development, neurodegenerative, and cancer pathways. These findings will help future mechanistic understanding of developmental disorders.

**IMPORTANCE:** First genome-wide BCL11b-protein and RNA interactome BCL11b interacts with RNA processing and splicing proteins BCL11b interacts with neurodegenerative genes and sarcoma genes BCL11b targets during cell proliferation and disease pathways

## INTRODUCTION

The B-cell lymphoma/leukemia 11B protein (Bcl11b, also known as COUP-TF-interacting protein 2, CTIP2) plays crucial roles in the epigenetic regulation of gene transcription and specifically in the control of the elongation process by interacting with an inactive form of the p-TEFb complex (7SK RNA, CDK9, cyclin T1, and HEXIM1) (Cherrier et al., 2013). BCL11b, which harbors six zinc finger domains (ZnFs) of C2H2-type, shares >60% homology with the human BCL11a, as well as the mouse, chicken, and Xenopus homologues. BCL11 proteins were isolated from T-cell lines derived from patients with T-cell leukemia, suggesting a role for BCL11 proteins in blood cell development and lymphomagenesis (Chiara et al., 2021; Fu et al., 2017; Helm et al., 2023; Montefiori et al., 2021; Sidwell and Rothenberg, 2021). Moreover, BCL11b was deleted in γ-ray induced mouse lymphomas, suggesting a role as a tumor suppressor and modulator of radiation-induced DNA damage, reviewed in (Kominami, 2012).

Studying the function of BCL11b by knocking out the gene is challenging (Longabaugh et al., 2017). Mice that lack the two alleles of BCL11b die shortly after birth, exhibiting defects in multiple tissues, including the immune system, central nervous system (CNS), skin, thymus, among other organs (Eto et al., 2022; Kominami, 2012; Przybylski et al., 2022; Simon et al., 2020), which suggests the importance of BCL11b during development. For example, in brain, BCL11b serves as a marker of layer V subcerebral projection neurons and striatal interneurons (Lennon et al., 2017, 2016; Simon et al., 2020). Several reports showed that BCL11b can regulate cell / tissue development, cell cycle, cell differentiation, and apoptosis in human, mouse, chicken and Drosophila, and multiple cell models, including cancer cells, blood cells, neuronal cells and mammary duct cells (Abe et al., 2023; Cai et al., 2017; Cherrier et al., 2009; Go et al., 2012; Grabarczyk et al., 2018; Liao et al., 2016; Miller et al., 2018; Parsa et al., 2023; Yang et al., 2020; Zhang et al., 2021; Zhao et al., 2017; Du et al., 2022; Holmes et al., 2021; Liau et al., 2023; Fox et al., 2022; Zhou et al., 2022). Although few interactions are known, the mechanisms by which BCL11b regulate cell proliferation and development and the target of BCL11b are largely unknown (Koumoundourou et al., 2024).

The absence of knockout BCL11b challenges the study of BCL11b functions as well as the interactions with protein and RNA in the cells. To our knowledge, BCL11b can form complex with 7sk ncRNA and p-TEFb complex (Cherrier et al., 2013), or with DNA-binding proteins, such as HP1 proteins (Rohr et al., 2003), NuRD (Cismasiu et al., 2008, 2005), HDACs (Marban et al., 2007), HMGA1 (Eilebrecht et al., 2014), DCAF1 (Forouzanfar et al., 2019), and KU proteins (Shadrina et al., 2020). The complexes including BCL11b and p-TEFb, HDACs, histone methylases, or transcription factors are essential to regulate HIV latency (Cherrier et al., 2013; Marban et al., 2007).

Although BCL11b has six C2H2 ZnF domains, to our knowledge, potential RNA-binding functions have not been well characterized for any of the BCL11 proteins nor the closely related proteins. In this study, we aim to identify BCL11b-binding complexes including those triggered by RNA, such as p-TEFb complex. Therefore, we used immunoprecipitation of proteins followed by mass spectrometry (IP-MS) as well as immunoprecipitation of cross-linked RNA (CLIP-seq), to identify the proteins and RNAs bound to BCL11b. Our results reveal that BCL11b can target RNA and proteins involved in neuronal development, cell cycle control, RNA splicing, cancers and neurodegenerative diseases. Our study offers new insights on role of BCL11b in diseases and open the route for future studies to understand the mechanisms of developmental disorders.

## MATERIAL AND METHODS

### Cell culture, plasmid, cell transfection and antibodies

The human microglial and HEK293T cell lines were maintained in Dulbecco’s modified Eagle medium (DMEM). The culture media were complemented with 10% (vol/vol) heat-inactivated fetal bovine serum (FBS) and 100 U/mL penicillin/streptomycin. The construct pFLAG-BCL11b and control pFLAG-pcDNA3 were previously used in (Cherrier et al., 2013). HEK293T cell cultured in 150-mm diameter dishes were transfected using a calcium phosphate co-precipitation method with the indicated plasmids. The whole cell lysates were prepared 48 hours post-transfection as previously described in (Cherrier et al., 2013). The following antibodies were used; BCL11b (A300–383A, Bethyl Laboratories), Goat anti-Rabbit IgG (HRP; 32460, Thermo Fisher), Drosha (ChIP Grade ab12286, Abcam), Mouse anti-IgG (CS200621, Millipore), Rabbit ani-IgG (PP64B, Upstate), Anti-flag M2 Affinity Gel (A2220, SIGMA), and FUS, UPF1 and m-IgGκ BP-HRP (sc-47711, sc-393594 and sc-516102, Santa Cruz).

### Protein extraction and co-immunoprecipitation (co-IP) for mass spectrometry

The cells seeded at a density of 3×10^6^ cells per dish and transfected with the indicated vectors (30 μg) using the method of co-precipitation with calcium phosphate. After 48 hours of transfection, the cells were lysed for 10 minutes on ice in a buffer containing: 10 mM HEPES, 1.5 mM MgCl2, 10 mM KCl, 0.5 mM DTT. After one minute centrifugation at 13,000 g, the nuclear pellet was lysed for 30 minutes on ice in another buffer containing: 20 mM HEPES, 25% glycerol, 1.5 mM MgCl2, 420 mM NaCl, 0.2 mM EDTA, 0.5 mM DTT. After 2 minutes centrifugation at 13000 g, the supernatant was collected and stored for further analysis by immunoprecipitation and western blot. Anti-proteases were systematically added to the lysis buffers. Immunoprecipitation was carried out on 500 μg of nuclear protein extracts by using the M2 anti-flag gel from SIGMA. The immunoprecipitated protein complexes were subjected to standard western blot analyzes. Proteins were detected using antibodies and visualized by a chemiluminescence detection system (Thermo Scientific Pierce ECL Western Blotting Substrate 32106).

### Mass spectrometry

HEK293T cells were transfected with the flag-BCL11b vector. The nuclear proteins were extracted and subjected to anti-flag immunoprecipitation. Immunoprecipitated proteins were eluted and then digested as described in this previous study (Turriziani et al., 2014). The peptides were separated and analyzed by reverse phase liquid chromatography coupled to a high-resolution mass spectrometer (Q-Exactive connected to an Ultimate Ultra3000 chromatography system (Thermo Scientific, Germany)). Mass spectra were analyzed using MaxQuant software. The results are representative of three independent experimental duplicates. The results were obtained in the form of protein intensities calculated by summarizing the ion intensities of the peptides which uniquely correspond to the protein sequence identified in all the samples (Turriziani et al., 2014). The proteins interacting with BCL11b were determined after normalization with the trypsin used for peptide digestion. Normalization with a control condition represented by cells transfected with the empty vector pcDNA3 has also been performed.

### Cross-Linking Immunoprecipitation Sequencing (CLIP-Seq)

The method has been described before (Maurin et al., 2015; Tabet et al., 2016). For short, cells were cultured and transfected with the pFLAG-BCL11b and pCDNA3 as control. The day prior the cell lysis the magnetic beads were prepared and incubated with lysis buffer only, with unspecific IgG, with anti-flag or with anti-BCL11b depending of the cell lysate to be analysed. Prior to lysis, cells were covered by ice cold PBS and exposed to UV light without for the crosslinking step by using the Stratalinker for 10 minutes at 4000J on ice. The cells were then lysed with lysis buffer (Tris HCl pH 7.4 50mM, CaCl2 0.08mM, SDS 0.1%, KCl 100 mM, MgCl2 2mM, NP40 1%, Deoxycholate 5g/L) complemented with RNasin (Promega) at 40U/ml and Proteinase Inhibitor Cocktail (Roche). 10 Units of TURBO-DNase (Ambion) was added for 5 minutes at 37 °C and the lysates were centrifuged for 15 minutes 13000 rpm 4°C. In the meantime, the tubes with the beads were washed with lysis buffer. Lysates were than subjected to two preclearing steps with beads alone and with IgG-beads to eliminate any unspecific binding before an overnight incubations with anti-flag, anti-BCL11b or unspecific IgG at 4°C. The day after the beads were washed three times with a high salt washing buffer (Tris HCl pH 7.4 50mM, EDTA 1mM, SDS 0.1%, KCl 1M, NP40 1%, Deoxycholate 5g/L) followed up by a Proteinase K treatment, 200μl per tube (100mM Tris HCl pH 7.4, 10mM EDTA, 40mM NaCl, Proteinase K 2mg/ml (Roche) for 20 minutes at 37° Celsius. The proteinase K activity was then stopped by adding 200μl PK Urea buffer (100mM Tris HCl pH 7.4, 10mM EDTA, 40mM NaCl, 7M Urea) at 37° C. The RNA extraction step followed up by adding 500μl of phenol/chloroform incubated 5 minutes at 37° C, 1100 rpm, the tubes were then centrifuged at RT at 13000rpm and the aqueous phase transferred to a new tube. A solution of 3M sodium acetate pH5.5 and 1μl Glycoblue was added with 1ml of 1:1 Ethanol:Isopropanol, the tubes were mixed and the RNA was left to precipitate overnight at -20° C. The tubes were then centrifuged at 13000rpm at 4 ° C for 10 minutes, the RNA pellet was washed twice with 500μl 80% ethanol and left to dry for 2 minutes with the lid open. The RNA was then dissolved in 10μl DPEC water and sent to sequence.

### RNA-Immunoprecipitation (RIP)

Microglial cells were cultured in 10cm dish, at 80%-90% confluence. The cells were lyzed and total RNA was extracted using NucleoBond Xtra Maxi (740414, Macherey-Nagel) according to the manufacturer’s recommendations. The lysate was separated from the cellular debris by centrifugation at 10000 rpm (10000g) at 4°C. The lysates were washed and pre-cleared for 1h at 4°C with 25μl Dynabeads protein G (Invitrogen) with unspecific IgG and incubated overnight at 4°C with 30μl Dynabeads previously coated with 5μg of the antibody of interest or unspecific IgG as negative control for overnight at 4°C. The RNP complexes were disrupted by performing an RNA extraction directly from the beads by adding 100μl chloroform, centrifuging at 13000 rpm (13000g) for 15 minutes. After this step, the liquid phase containing the RNA was separated, mixed with isopropanol and Glycoblue (Invitrogen) to facilitate the formation of the RNA pellet during 10 minutes incubation on ice and followed by centrifugation at 13000 rpm at 4°C. The pellet was washed with ethanol 80% and re-suspended in 16μl of DEPC water. RNAs were reverse transcribed to cDNA using iScript Reverse Transcription Supermix for RT-qPCR (BIO-RAD) in a final volume of 20ul. cDNA was then diluted 5 times and 2μL are used for the qPCR analysis with SsoAdvanced Universal SYBR Green Supermix (BIO-RAD).

### Protein extraction and co-immunoprecipitation for western-blotting

HEK293T cells were transfected with Flag-BCL11b and control plasmids with JetPei transfection reagent. After 48 hours, proteins were extracted with a RIPA buffer (150 mM 186 NaCl, 1% Nonidet P-40, 0.1% SDS (10%), 1.5mM MgCl2, 50 mM Tris-HCl, pH 8). 4 µg of anti-flag or anti-IgG control antibodies were incubated with Dynabeads protein G in low salt IPLS buffer (50 mM Tris HCL PH 7.5, 120 mM NaCL, 0.5 mM EDTA, 0.5 % Nonidet P-40, 10 % Glycerol) and washed before addition of 1mg of protein extracts and incubated overnight at 4°C. The IP samples were washed twice with 1 ml of IPLS buffer at 4°C, then twice with high salt concentration buffer IPHS (50 mM Tris HCL PH 7.5, 0.5 M NaCL, 0.5 mM EDTA, 0.5 % Nonidet P-40, 10 % Glycerol) and finally twice with IPLS. EDTA-free Protease Inhibitor Cocktail (Roche) was added to the IPLS and IPHS buffers used for the washing in order to avoid protein degradation. SDS PAGE gel electrophoresis was performed and the proteins were transferred to a nitrocellulose membrane using the Trans-Blot Turbo Transfer system from Bio-Rad. After the transfer of proteins, the membranes were incubated for 5 minutes in the blocking buffer from Bio-Rad (Every-blot blocking buffer, Ref#12010020). After the blocking step, the membranes were incubated with primary and secondary antibodies mentioned in Annex A (The antibodies dilutions were made in the Every-blot blocking buffer). After immunoblotting, the detection was carried out using the Super Signal Chemiluminescence Detection System (Thermo Fisher), and were visualized by Fusion imaging system (Vilber Lourm).

### PCR and RNA quantification

The total RNA was extracted using NucleoBond Xtra Maxi (740414, Macherey-Nagel) according to the manufacturer’s recommendations. RNA was reverse transcribed into cDNA using iScript Reverse Transcriptase Supermix (BIO-RAD) and oligo 12–18 Primers (Thermo Fisher). The quantitative PCR was performed on the diluted cDNA (in DNase-free water) using SsoAdvanced Universal SYBR Green Supermix (BIO-RAD). The PCR products were then run into gel to confirm the size. The primers pairs used in the study are listed in table S4. Additional PCR test were performed to confirm non-genomic DNA contaminations.

### Mass spectrometry bioinformatics analysis

Proteins with three or more MS signals (MS/MS Count) were selected. The proteins should have four or more peptides and at least one unique peptide to be selected. In case of two or three peptides with at least two razor + unique peptides were selected, on condition that the MS/MS count > 3. The unique sequence coverage was taken into consideration (over 1%). In all cases, the p-values between the condition and control were calculated using student t-test. Proteins with p-values < 0.05, and two-fold increase (1 log2 fold increase) were selected. For downstream GO analysis, we used combination of methods, including the GO construction (http://geneontology.org/) and DAVID database (https://david.ncifcrf.gov/). Additionally, we downloaded OBO gene annotation files for proteins and ncRNA from (http://geneontology.org/docs/download-ontology/) in 2018. For protein complexes, we obtained the data from CORUM database (http://mips.helmholtz-muenchen.de/corum/).

### CLIP sequencing and bioinformatics analysis

After CLIP-seq experiments, the RNA library was sequenced on Illumina Hiseq 4000 sequencer as Single-Read 50 base reads following Illumina’s instructions. FastQC (https://www.bioinformatics.babraham.ac.uk/projects/fastqc/) was run to produce base quality.

After adapters were cut, the short reads less than 10 bases were removed. Reads were mapped onto the *Homo sapiens* genome (hg19 assembly) using Bowtie2 (http://bowtie-bio.sourceforge.net/bowtie2/), Samtools (http://samtools.sourceforge.net/), or STAR aligner (https://github.com/alexdobin/STAR). The number of reads per gene, transcript or exon counted using featureCounts, Subread package (http://subread.sourceforge.net/). The genome assembly file was downloaded from GENCODE (https://www.gencodegenes.org/). The protein-coding genes, exons, TSS and transcript annotation file was downloaded from UCSC (http://hgdownload.cse.ucsc.edu/) or Ensembl (https://grch37.ensembl.org/), RNA annotation from (https://rnacentral.org/), and OBO gene from (http://geneontology.org/) in 2018. The CLIP-seq reads are visualized using Integrative Genomics Viewer (IGV) software (https://igv.org/).

The three control conditions and three replicates in both HEK cells and microglial were used for the analyses. The uniquely mapped reads (MAPQ=255) were selected. This leads to on average 1.1 or 2.3 for million unique hits from control and replicate HEK experiments, respectively, and 1.1 or 2.1 million unique hits from control and replicate of microglial experiments, respectively. Then average of RNA CLIP tags (reads) in control conditions and replicates were calculated for whole human genome. In case of microglial, the top binding genes should pass three cut-offs, RNA CLIP tags over 4-fold (log2=2) to the control, the number of tags of experiments should be more than 1000 those in control cases (average of number of tags of experiments – average of control ≥1000), and the p-values ≤ 0.05 cut-offs. For HEK cells, two-fold the control (log2 fold change =1), the average of number of tags of experiments – average of control ≥200, and p-values ≤ 0.1. We selected the genes that passed the cut-offs in both microglial and HEK cells datasets at the same time. This leads to a final dataset of 740 genes, which encode RNA transcripts bind to BCL11b as shown by CLIP-seq method. The dataset then used for further GO analysis and other bioinformatics analyses.

To quantify the amount of RNA CLIP reads per transcripts, exons, and introns features, subread package was used. For this step un-filtered BAM files were used, i.e., the reads can be mapped to multiple features. Number of exons and introns were obtained from Ensembl BioMart or UCSC refGene file. To get insights into the differential binding between BCL11b and the different transcripts encoded from the same gene, we chose 23 genes. These genes are enriched with significant CLIP reads and they characterize by the encoding both protein-coding and non-coding transcripts. Boxplots were plotted using R BoxPlotR tool (http://shiny.chemgrid.org/boxplotr/). Genes were visualized by UCSC browser (https://genome.ucsc.edu/) or IGV (http://software.broadinstitute.org/software/igv/).

For over-expressed clusters, the significant cluster is two-fold the control. As a second method, we counted the number of RNAs per 10kb-long bins of human genome. The clusters that pass the threshold of 5-fold to the control were selected, leading to 646 cluster. The default parameters were selected. We then selected the clusters that experiments are enriched over control. This leads to results of 1466 and 1583 clusters in HEK293T and microglial, respectively. In all cases, the significant clusters were selected for gene ontology and disease ontology using DAVID, GO construction and GREAT tool (http://great.stanford.edu/). The R packages “ClusterProfiler, EnrichGO, and Enrichplot” are used to determine the enrichment scores. The “Pathview” R package is used to construct KEGG pathways. Volcano plots are constructed using “EnhancedVolcano” R package.

## RESULTS

### BCL11b is associated with cellular RNA

Although BCL11b has six C2H2 ZnF domains, to our knowledge, potential RNA-binding functions have not been well characterized for any of the BCL11 proteins; except BCL11b can bind the 7SK RNA to regulate transcription elongation (Cherrier et al., 2013; Eilebrecht et al., 2014). To examine the RNA-binding activity and obtain a transcriptome-wide landscape of BCL11b-bound RNAs, we cross-linked BCL11b to cellular RNAs using UV radiation, followed by immunoprecipitation and sequencing (CLIP-seq). These experiments were performed by targeting endogenous BCL11b in microglial cells, or Flag-BCL11b in HEK293 cells. Microglia were previously used as model cells to study roles of BCL11b on gene silencing and HIV latency (Cherrier et al., 2013; Desplats et al., 2013; Eilebrecht et al., 2014). The top genes that passed the cut-offs (see material and methods) in each cell line were only selected which led to a dataset of 740 different genes that have high CLIP-seq reads in both HEK and microglial cells (Figure S1, S2, Suppl. data S1), we call this dataset as BCL11b-CLIP-gene-set. Interestingly, we found that over 90% of RNAs in this BCL11b-CLIP-gene-set are mapped to protein-coding genes (Figure S1, Table S1, and Suppl. data S1). GO terms of the BCL11b-CLIP-gene-set include genes belong to development of the brain, neuron, hemopoiesis, blood vessels, and hair follicles (details are found in Figure S3, Suppl. data S2). Additionally, BCL11b binds to RNA transcribed from genes involved in RNA processing, for example, BCL11b binds to protein-coding RNAs and non-coding RNAs (ncRNAs), involved in biogenesis of the major and the minor spliceosome (Figure S1, Table S2, Suppl. data 2).

The pathway analysis of BCL11b-CLIP-gene-set reveals regulation of RNA transcription of multiple immune response pathways including Wnt signaling and MAPK signaling (Figure 1A, Figure S2, Suppl. data 2). BCL11b can regulate the same genes in Wnt signaling in different cell lines, including microglia and HEK cells (Figure 1B, S3, S4, Suppl. data 1). Noteworthy, we collected list of genes that harbor the top BCL11b CLIP-seq reads and compared with previously published differential expression transcriptomics data (by microarray and RNA-seq analyses), we observe that our study identifies novel genes and pathways that have not been previously described (Figure 1B, S4) (Cherrier et al., 2013; Drashansky et al., 2021, 2019; Eilebrecht et al., 2014; Hasan et al., 2019; Hosokawa et al., 2018; Longabaugh et al., 2017; Tang et al., 2011). Since BCL11b functions on gene transcription have been previously reported, these results suggest that BCL11b may also bind some RNA to regulate their processing. Surprisingly, we found CTIP2 bound to RNA coding for regulators of the RNA processing and RNA splicing pathways, such as EWSR1, FUS, SFPQ, SRRM2, and DNMT1, in addition to ncRNA (Figure 1C, and figures S1), specifically to exon and exon-intron junction regions (Figure 1D, E, Figure S5-S6). Noteworthy, we could not observe significant changes in the expression of CTIP2-bound protein coding RNA upon over-expression of BCL11b (Figure 1E).

**FIGURE 1.**
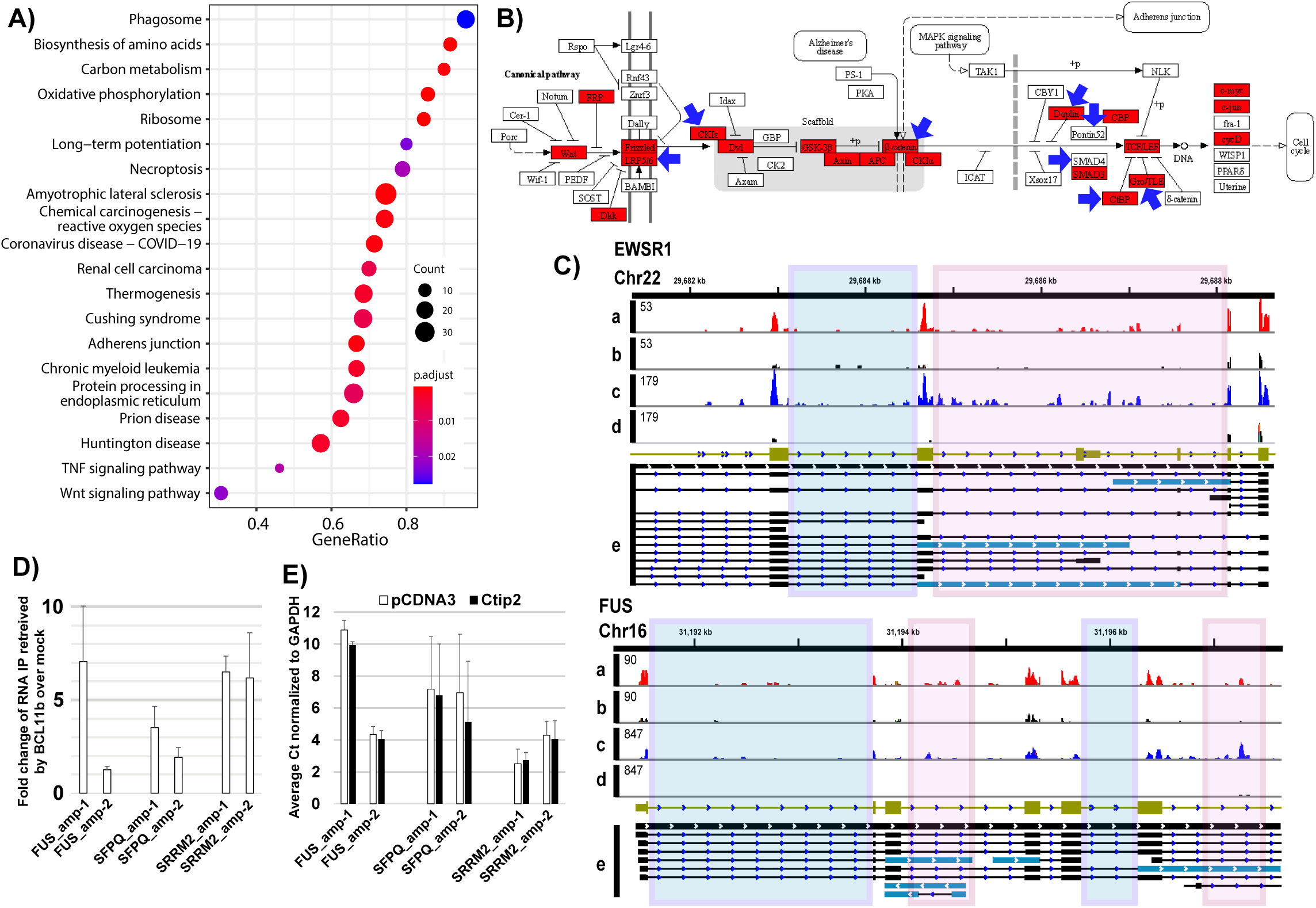
BCL11b-RNA interactions as deciphered by CLIP-seq. A) KEGG pathways of genes encode RNA bound to BCL11b, Figure S1. B) Wnt signaling is among pathways that are regulated by BCL11b-RNA interactions, the red box means the CLIP reads increased by at least 4 log2 fold change compared to control in microglia, blue arrow means CLIP reads increased at least 2 log2 fold after over-expression in HEK cells, for full pathway Figure S2. The GSEA and KEGG pathway were constructed using “ClusterProfiler, EnrichGO, Enrichplot, and Pathview” R packages. C) Representative CLIP-seq results of three protein-coding genes (EWSR1 and FUS), visualized using Integrative Genomics Viewer (IGV) tool. The olive-green bars refer to the NCBI RefSeq genes. X-axis represents the coordinates of the gene and the chromosome number. The Y-axis represents the number of CLIP reads in each condition. a) Over-expressed BCL11b in HEK cells, b) endogenous BCL11b in microglial cells, whereas, c and d are their control experiments, respectively. e) The black bars represent the GENCODE transcripts of the stated genes, the sky-blue transcripts refer to the protein non-coding transcripts that harbor significant CLIP reads and their coordinates correspond to intron of RefSeq gene, these intronic regions are highlighted with light rose colored boxes. The light blue horizontal boxes refer to introns without significant BCL11b CLIP reads. For additional figures and CLIP-seq reads alignments, see Figure S1. D) Results of RNA-IP-qPCR (RIP-PCR) of three genes, the fold change of RNA bound to endogenous BCL11b to the input control, and E) gene expression of the amplicons used in D using overexpressed BCL11b plasmid and pCDNA3 vector. For PCR primers and raw Ct values, Table S4-S5, and locations of the amplicons and gel amplification, Figure S4, S5.

To confirm these results, we divided the whole genome into 10kb regions (bins) and counted the number of reads corresponding to each bin (see material and methods section). For additional confirmation, we performed peak calling to cluster the RNAs into bins. We then selected the regions that were enriched over the control replicates of the two cell lines. The results showed that over 40% of CLIP reads mapped within 0-50kb down-stream of the transcription start sites (TSS), and about one third up-stream to TSS (Figure S7-S8, Suppl. data S3). Gene ontology analysis for the genes localized in these regions suggested their involvement in angiogenesis, response to stress, and inflammation pathways, whereas disease ontology shows involvement in cancers, Alzheimer, and Huntington diseases (Figure S7-S8, Suppl. data S3).

Taken together, our results demonstrate that half of the BCL11b-binding RNAs are mapped to gene body regions, suggesting that BCL11b may regulate co-transcriptional RNA processing and/or mRNA biogenesis.

### BCL11b binds to multiple RNA isoforms of the same genes

To further examine the association of BCL11b with RNA transcripts (isoforms), we counted the number of reads per exon for CLIP reads that mapped to genes. We found that about 75% of the CLIP reads mapped to exons (Figure 2A). Analysis of the intronic reads revealed that about 25% of the reads corresponded to isoforms that will not yield stable protein products (we refer to them here as isoforms that does not code for proteins, INCP). For example, number of BCL11b CLIP-seq reads are mapped to intronic regions within EWSR1, FUS, and SRRM2 genes, which code for retained introns (IR), nonsense mediated decay (NMD), non-stop decay or processed transcript, (Figure 1C, Suppl. Figure S1). This result is consistent with our proteomics data (shown in the further section) demonstrating that BCL11b interacts with proteins involved in RNA splicing and NMD pathways, including FUS, SMN1, RNPs, Drosha, and UPF1.

**FIGURE 2.**
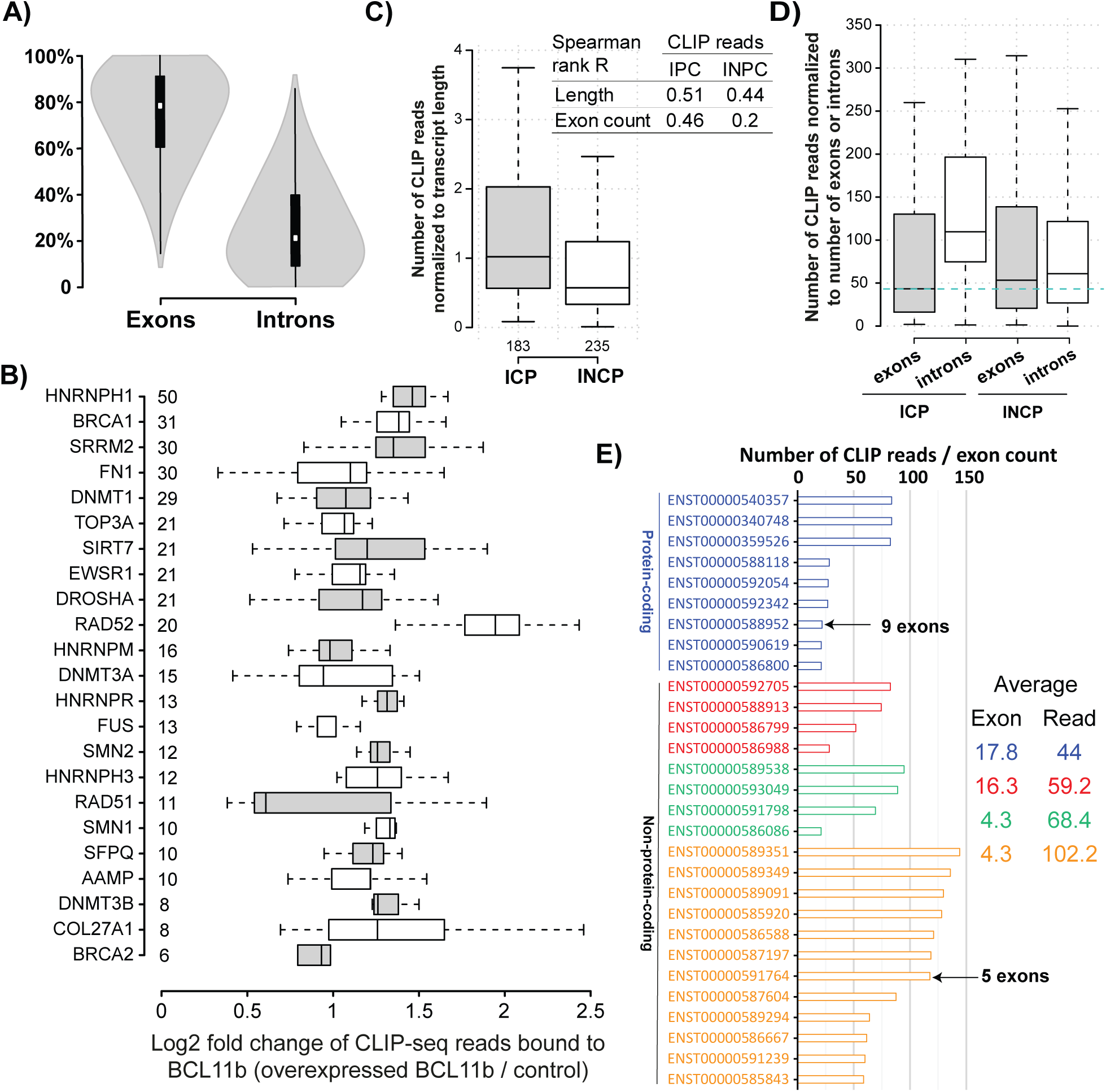
BCL11b-bound RNA at transcript (isoform) level. A) Violin shape represents the number of CLIP reads mapped to exons and introns of the significant genes. B) BCL11b binds differentially to the selected 23 genes. Number next to the names are number of transcripts. C) Numbers of CLIP reads normalized to number of exons for isoforms coding for proteins (for short, ICPs) or isoforms do not code for proteins (for short, INCPs). The table shows the Spearman’s rank correlation coefficient (R) between numbers of CLIP reads in case of ICPs or INCPs from one side and length or number of exons from the other side. D) Number of CLIP reads mapped to exons and those mapped to introns in case of ICPs or INCPs. E) Number of CLIP reads mapped to exons of DNMT1 gene are counted and normalized to number of exons. The table shows the average number of CLIP reads and number of exons within protein-coding transcripts and non-protein-coding transcripts. The blue colors refer to ICPs, whereas red, green and orange colors refer to nonsense mediated decay, processed transcript, and retained intron transcripts, respectively. Figures 3B-3E were constructed from Suppl. data S4.

To further test this finding, we collected the transcripts (or RNA isoforms) of 23 genes that encode isoforms coding for proteins (ICPs) and isoforms that do not code for proteins (INCPs), the latter would be indicative of NMD or IR transcripts / isoforms. We then counted the number of reads that correspond to each isoform of the gene, including the protein-coding isoforms (ICPs) or those not coding for proteins (INCPs). We found that BCL11b binds to different isoforms of the same gene in differential manner (Figure 2B, Suppl. data S4). For example, the TOP3A gene encodes 21 different RNA isoforms, similarly ESWR1, Drosha, SIRT7 and RAD52 genes encode 20-21 RNA isoforms. The fold increase to control of CLIP-seq for TOP3A is about 1.2-folds, whereas the binding to RNA of RAD52 is up to 7-folds increase (Figure 2B). The same concept applies to SMN1 (10 isoforms), COL27A1 (8 isoforms) and RAD51 (11 isoforms). We counted the CLIP-seq reads per each transcript (or isoform) and then normalized to the length of the transcripts. BCL11b favors binding to long isoforms coding for protein (ICPs) than protein non coding (INCPs), Spearman correlation rank (R) between the CLIP reads and transcripts length for ICPs is 0.51 and 0.44 for INCP, which suggest that transcript length could be a contributing but not a major factor (Figure 2C). Together these results suggest that BCL11b can distinguish between different RNA isoforms of the same gene, which is crucial function for a protein involved in RNA metabolism or cell development.

We still noticed some CLIP reads that correspond to intronic regions within ICPs, but the CLIP reads are mapped to the coordinates of exons in other INCP isoforms. Noteworthy, NMD or IR transcripts introduce additional exons instead of introns. To explain this concept, we highlighted the genomics introns in pink colors, see (Figure 1C and S1). Although these intron regions are usually removed in case of ICP transcripts (thick dark bars in the panel e of Figure 1C and S1), we found that part of these regions corresponded to exons in INCPs (light blue bars in e panel in Figure 1C and S1). Computationally, we counted number of reads mapped to exons and introns, we normalized them to the number of exons or introns as shown in Figure 2D and Suppl. data S4. If BCL11b binds to exons regions of RNA, we expect to find that most of the CLIP reads are biased to exons coordinates. However, in Figure 2D, we observed that most of the reads are mapped to introns. In case of INCPs, the number of reads were almost equally distributed throughout the exons and introns. Statistically, we calculated the Spearman correlation rank (R) between number of reads and number of exons, which show that number of exons correlates with number of CLIP reads, in case of ICPs, but not in case of INCPs (R of INCPs=0.2, and R of ICPs=0.46). This result suggests that INCPs with few numbers of exons have multiple CLIP reads. The DNA (cytosine-5-)-methyltransferase 1 (DNMT1) transcripts are good examples that support this observation. Number of reads mapped to retained introns (IR) IR transcripts are 2-3-fold higher than the number of reads mapped to INCPs transcripts. Indeed, the 5 exons IR transcripts (ENST00000591764, with 589 reads) have 3-folds more reads than the 9 exons protein coding transcripts (ENST00000588952, with 196 reads) (Figure 2E, Suppl. data S4).

As mentioned above, comparing top genes that harbor highest CLIP-reads with publicly available transcriptomics data showed that most of the genes are not differentially expressed after BCL11b loss or gain of function (Cherrier et al., 2013; Drashansky et al., 2021, 2019; Eilebrecht et al., 2014; Hasan et al., 2019; Hosokawa et al., 2018; Longabaugh et al., 2017; Tang et al., 2011).

Taken together, our CLIP-seq results show that BCL11b can recognize the different type of RNA isoforms (either protein coding or not coding), and binds to the isoform in RNA type-specific manner. This characteristic is very important for the proteins that play roles in selecting the correct isoforms to be translated (i.e. isoform selection). On other words, this finding suggests that BCL11b may bind to specific RNA isoforms to regulate RNA processing or metabolism at post-transcriptional levels. However, experimental confirmations will be needed to confirm this hypothesis.

### Quantitative protein interactomics of BCL11b

To characterize the protein interactants and pathways triggered by BCL11b, we used first HEK cells expressing Flag-BCL11b. After isolation of BCL11b and associated proteins using anti-flag antibodies, we performed quantitative LC-MS/MS. After statistical analysis, we identified 629 BCL11b protein partners (>two-fold change and p-value <0.05; Figure S9, supplementary dataset S5).

#### BCL11b interacts with DNA damage repair proteins and transcription regulators (e.g. HIV-1 transcription)

Our proteomics analysis shows that BCL11b binds to chromatin organization, and DNA recombination proteins, including HP1 and DCAF1 proteins, and components of NuRD and SNW1 complexes. Additionally, BCL11b binds to protein regulators of DNA damage repair, such as H2AX, BRCA2, MCMs, PARP1, and SMCs proteins. BCL11b regulate the transcription through interactions with multiple epigenetics and transcription regulators including CCNT1, CDK9, SMARCAs and SMARCCs transcription regulators, as well as histone methyltransferases and acetyltransferases (KMT2A and HDACs). In agreement with our results, previous publications from our lab and colleagues reported that over-expression of BCL11b in HEK cells and other cell lines shows similar interactions, including HDACs, HP1, HMGA1, DCAF1, KU proteins, NuRD complex, and P-TEFb complex (CDK9 and CycT1) (Cherrier et al., 2013; Cismasiu et al., 2008, 2005; Eilebrecht et al., 2014; Forouzanfar et al., 2019; Marban et al., 2007; Rohr et al., 2003; Shadrina et al., 2020; Vickridge et al., 2024) (Figure 3A, S10 and suppl. data S5). The interactions between BCL11b and the transcription regulators were shown to be essential during HIV-1 latency, or epigenetics modifications and DNA damage repair mechanisms. Noteworthy, some of these interactions can be dissociated after RNase treatment, such as BCL11b-P-TEFb interaction, but not BCL11b-HDAC, which means that some of these complexes are RNA-dependent (Cherrier et al., 2013). Noting that we detected a hit of ribonucleases in our MS results, the role of BCL11b in these ribonucleoprotein complexes deserves to be studied by future independent studies.

**FIGURE 3.**
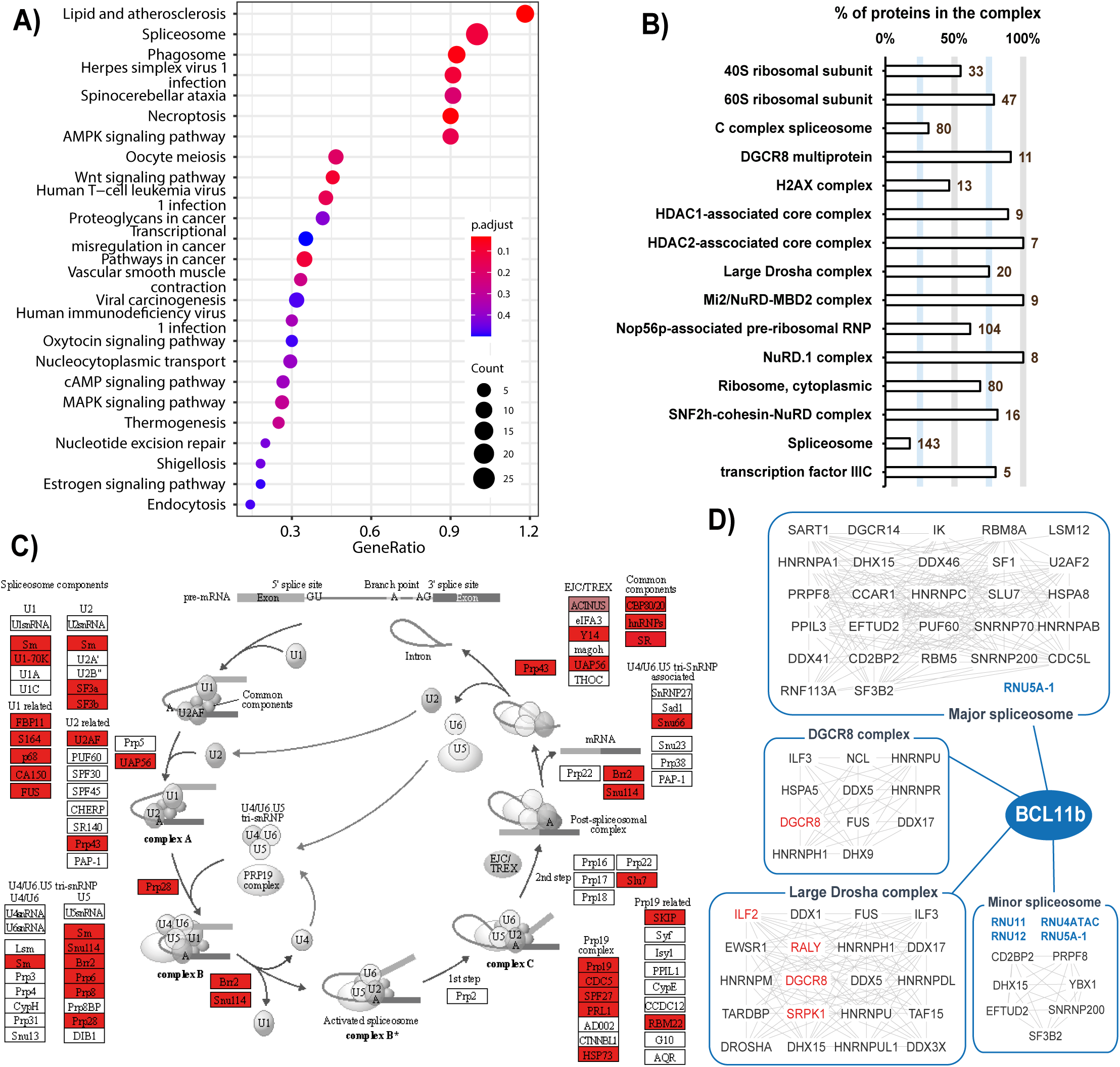
BCL11b-interacting proteins (for short BIPs) as deciphered by IP-MS. A) KEGG pathways of BCL11b-interaction proteins, additional details on GO terms and GSEA analysis can be found in Figure S9, SI data S5. B) The complexes that constitute of BIPs, X-axis show percent of BIPs to the total number of proteins in the complex; the total number of proteins in the complex is written in dark blue outside the bars. Complexes were downloaded from CORUM database, SI data S5. C) KEGG spliceosome pathway (KEGG ID: hsa03040), the red boxes represent BIPs, for additional KEGG pathways, Figure S10-S11, SI data S5. D) Four representative complexes with the name of proteins. The BIPs are found in rectangles, BCL11b-interacting ncRNAs are in blue, whereas the non-BIPs are in red colors. The arrows refer to protein-protein interactions. The interactions network were downloaded from STRING database and constructed using Cytoscape software. For additional complexes and interactions, Table, S2-S3, SI data S5.

#### BCL11b interacts with proteins dedicated to cell cycle and epigenetic regulations

We observed that BCL11b interacts with proteins and complexes that are involved in DNA replication and cell cycle (Figure 3A, S10, S11 and supplementary data S5), including CDK proteins. Cyclin-dependent kinases (CDK) are group of proteins that regulate cell cycle and transcription by phosphorylating their substrates. CDK1, 2, 4 and 6 among other were found to bind to BCL11b (Figure S11). In agreement with previous reports that showed BCL11b can regulate cell cycle, cell differentiation, and apoptosis by interacting with long non-coding RNAs and impairs p21, p16, cyclin D1 and CDK2 pathways (Abe et al., 2023; Cai et al., 2017; Cherrier et al., 2009; Go et al., 2012; Liao et al., 2016; Parsa et al., 2023; Zhang et al., 2021; Zhao et al., 2017). However, the exact mechanism by which BCL11b impairs cell proliferation is not known. Our data highlight the interactions between BCL11b with multiple cyclin dependent kinases (e.g. CDK1, 2, and 4/6), cell division control protein (e.g. CDC7, CDC20, and CDC45), and minichromosome maintenance complex (MCM) proteins, in addition to SKP1 a member of the SCF ubiquitin ligase protein complex.

#### BCL11b interacts with proteins involved in translation and RNA splicing

Among the novel findings of our study is the interactions between BCL11b with multiple proteins involved in RNA processing and translation, such as 40S and 60S ribosomal subunits (Figure 3A-D, S10, S12, supplementary data S5). Moreover, BCL11b interacts with many RNA-binding proteins, including FUS, spliceosomal subunits, small nuclear RNP (snRNPs), heterogeneous nuclear RNPs (hnRNP), serine/arginine (SR)-rich proteins and the survival motor neuron (SMN) protein complex. Overall, BCL11b interacts with 22 different hnRNPs or snRNPs, and about 17% of proteins that constitute the spliceosome (Table S2, S3, Figure S10, S12, suppl data S5).

FUS is another crucial factor that is involved in RNA splicing, RNA transport, and regulation of gene expression in response to DNA damage (Chen et al., 2019). FUS can accumulate near the transcription start sites (TSS) and binds to transcription initiation factors to inhibit phosphorylation of the RNA polymerase 2 C-terminal domain (POLR2-CTD) (Chen et al., 2019). We performed co-IP and immunoblot experiments to confirm BCL11b interactions with SMN1 and FUS (Figure 4A, 4B). Taken together, our IP-MS and IP-immunoblot experiments establish that BCL11b interacts with SMN1 and FUS proteins, suggesting that BCL11b may contribute to RNA splicing and/or translation. However, further investigations will be needed to confirm BCL11b functions on RNA processing.

**FIGURE 4.**
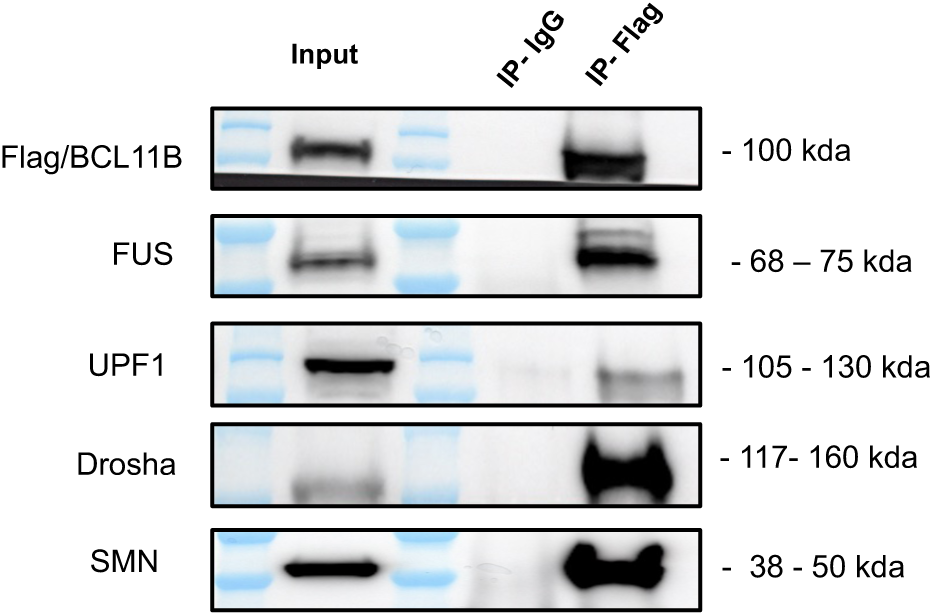
BCL11B interacts with FUS, Drosha, SMN1 and UPF1. Coimmunoprecipitations targeting Flag-BCL11B were performed, and the indicated partners were detected by Western blot with the corresponding antibodies.

#### BCL11b interacts with the Drosha and the DGCR8 protein complexes

FUS is a member of the Drosha and the DGCR8 complexes, which are able to regulate gene expression at transcriptional and post-transcriptional levels (Pong and Gullerova, 2018). On the other hand, Ago2 is the key protein in the RNA interference (RNAi) pathway. Ago2 can bind to Dicer or TRBP proteins, and Drosha or TRBP complexes can regulate mRNA stability or translation (Pong and Gullerova, 2018). Our IP-MS data revealed that BCL11b is associated with 15 of the 20 Drosha complex proteins and 10 of the 11 DGCR8 complex proteins (Figure 3D, S10, S12). In addition, 20 proteins of TRBP complex, including Ago2, were observed associated with BCL11b (supplementary dataset S5). To confirm these results, we performed co-IP experiments, which verified the association of BCL11b with FUS and Drosha (Figure 1A, 1B and 1C), further suggesting that BCL11b may coordinate with FUS and Drosha to regulate gene expression at transcriptional and post-transcriptional levels, such as mRNA translation and stability.

#### BCL11b binds to protein from the nonsense-mediated mRNA decay (NMD) pathway

The NMD pathway degrades RNA transcripts with premature stop codons (Kurosaki et al., 2019). UPF1 is a key regulator of the NMD pathway. Among the significant IP-MS hits, we observed interactions between BCL11b with proteins involved in the NMD pathway. Out of 120 proteins identified in this specific GO molecular pathway, 61 were identified in the BCL11b data set (Figure 3B, 3C, S10, S12), among these proteins two core NMD proteins (SMG7 and UPF1), along with one member of the exon junction complex (RBM8A), as well as multiple ribosomal proteins. It is worth to mention that UPF1 can be recruited during mRNA nuclear export, ribosome stalling, or after mRNA translation termination. Co-IP experiments confirmed that BCL11b associates with UPF1, linking BCL11b with proteins that control NMD (Figure 4D).

### BCL11b interacts with RNA transcripts and proteins from the same gene

We found BCL11b associated with 137 RNA transcripts and proteins encoded by the same gene (Figure 5A, and Suppl. data S6), including EWSR1, FUS, SPFQ, UPF1, tubulins, CHDs and hnRNPs. Interestingly, as we described earlier, BCL11b binds to multiple proteins and ncRNAs constituents of spliceosomes (Table S2, S3, Suppl. data S6). GO analysis revealed that these proteins are involved in RNA processing, NMD, and translation (Figure 5B, Suppl. data S6).

**FIGURE 5.**
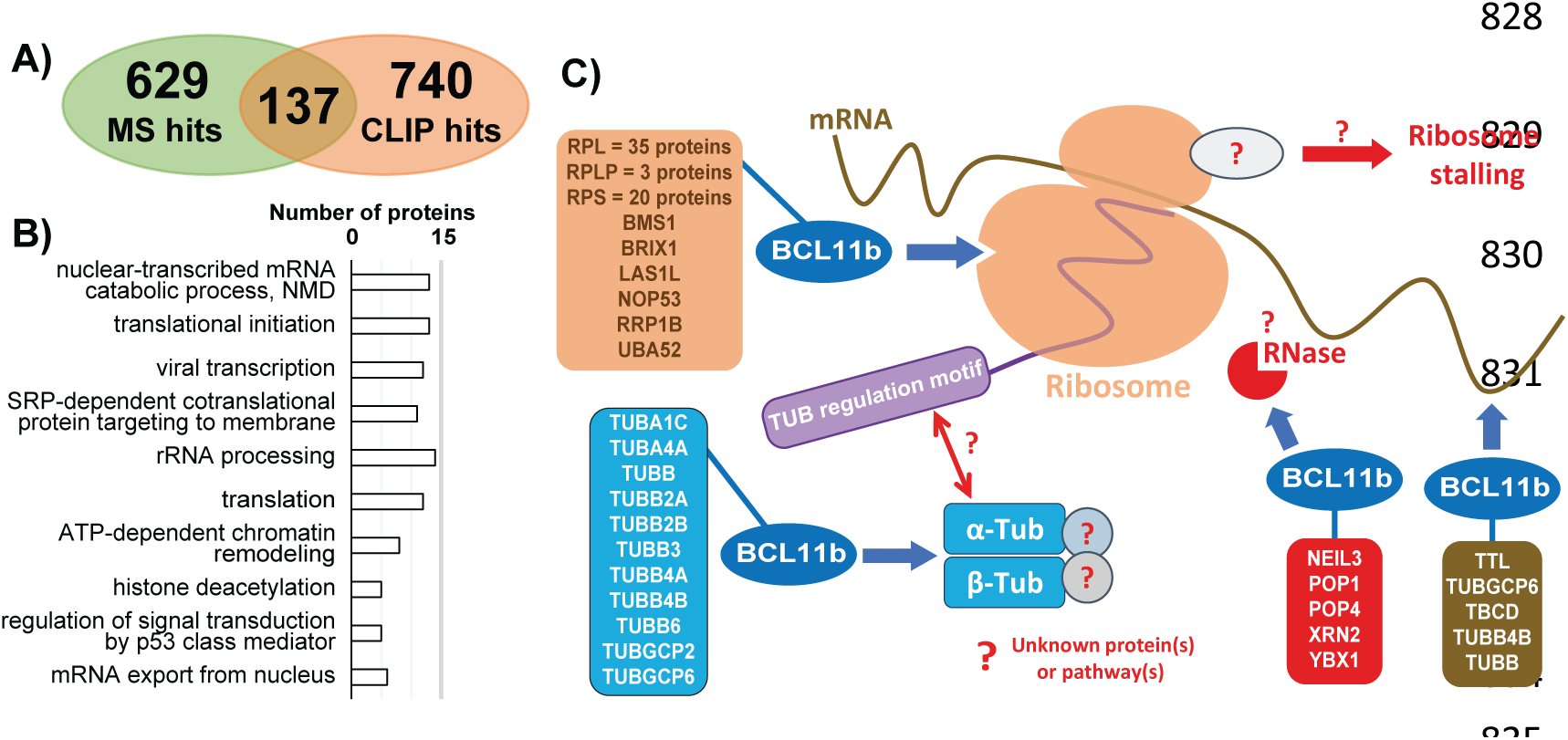
BCL11b binds to RNA and proteins encoded from the same genes. A) Number of BCL11b-interacting proteins, from IP-MS experiment, that passed our cut-off (so-called MS hits); number of genes, which have CLIP reads passed our cut-off (so-called CLIP hits); 137 genes can bind to BCL11b in form of RNA or in form of protein. B) GO terms of 137 shared genes (BCL11b binds to RNA and protein product). C) Schematic diagram of the possible tubulin autoregulation mechanism. BCL11b binds to multiple tubulin RNA, in brown box, in addition BCL11b binds to multiple ribosomes (oranges box), tubulins (blue box) and ribonucleases (red box).

To our knowledge, the only known case of a protein that binds to its RNA is a tubulin. For short, tubulin was reported to use a unique and unknown mechanism to auto-regulate its expression level within the cell to ensure a stable production of the protein in cells, and over-expression of tubulin gene does not lead to increase its protein expression level. It is thought that tubulins α and β could polymerize through an unknown mediator, which then binds to the nascent tubulin peptide, ribosomes and/or ribonucleases to terminate the translation (Gasic and Mitchison, 2019). We found that BCL11b interacts with multiple isoforms of tubulin proteins and tubulin RNAs, as well as ribonucleases and ribosomes, i.e. multiple members of the same protein family (Figure 5C). Noteworthy, RNPs and ribosomal proteins are assembled in nucleus before translocalized to cytoplasm, whereas tubulins are required for spindle formation during cell cycle phases. Again, these results support the involvement of BCL11b in the regulation of protein expressions at post-transcriptional levels, particularly the genes involved in RNA splicing and cell cycle.

### Involvement of BCL11b-associated factors in diseases

Our MS and CLIP-seq results show that BCL11b interacts with proteins that involved in cancer, Ewing Sarcoma, or neurodegenerative diseases (Figure 6, S13-S14 Table S2-S3), such as EWSR1, SMN1, FUS, TDP-43 and TAF15, SRRM2, as well as ncRNAs, such as 7SK, MALAT1, NEAT1. We then searched bibliography and public databases to confirm our finding related to the roles of BCL11b in disease. We found that BCL11b is modulated in lymphoma and Leukemia (Fang et al., 2021; Fu et al., 2017; Kominami, 2012; Montefiori et al., 2021), Ewing Sarcoma (Orth et al., 2020; Wiles et al., 2013), as well as in neuron diseases, such as motor neuron amyotrophic lateral sclerosis (ALS), spinal muscular atrophy (SMA), Parkinson’s disease (PD), and Huntington’s disease (Ahmed et al., 2015; Doktor et al., 2016; Harb et al., 2022; Lennon et al., 2017, 2016; Maeda et al., 2014; Scherzer et al., 2007; Srakočić et al., 2023), summarized in Figure 6A. However, the role of BCL11b in these diseases was not clear, because of the lack or the difficulties of the functional studies related to protein and RNA interactions of BCL11b.

**FIGURE 6.**
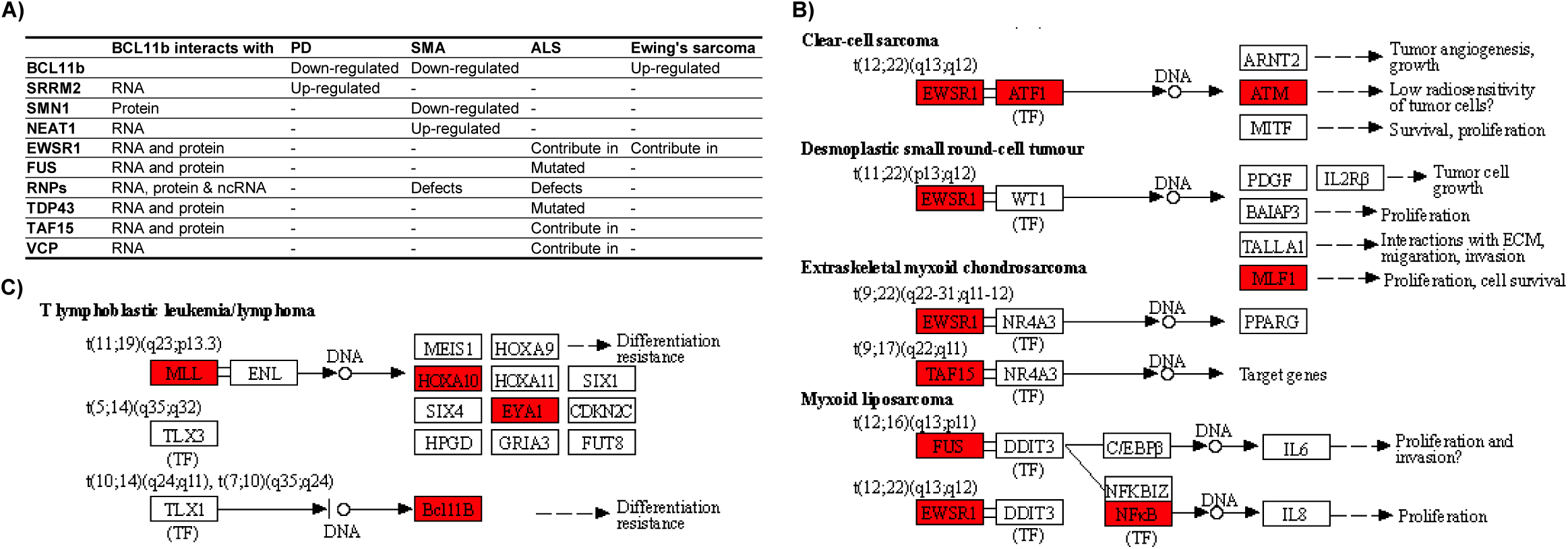
BCL11b is involved in multiple diseases. A) The table shows the contribution of BCL11b and other genes in the Parkinson’s disease (PD), SMA, ALS and Ewing sarcoma diseases. B and C) are examples of BCL11b-interactions proteins contributing to transcriptional mis-regulation in cancer, sarcoma, and lymphoma pathways (KEGG ID: hsa05202), for additional KEGG disease pathways, Figures S12-S13.

## DISCUSSION AND CONCLUSION

This study describes for the first time BCL11b-associated protein and RNA interactome (Figure 7). To our knowledge this is the first time to report a protein to bind to RNA transcript and protein encoded from the same gene, such as FUS. We showed that BCL11b binds to RNA and proteins encoded from genes regulating development, cell cycle, transcription, and translation. In addition to its function as a transcription regulator, BCL11b binds to ribonucleic complexes such as ribosomes and RNPs. We confirmed that BCL11b binds to proteins involved in RNA processing, such as RNA splicing and NMD pathway, in addition to the ncRNA that are involved in RNA splicing. First, we demonstrate that BCL11b binds to RNA processing proteins (such as FUS, Drosha, Ago2, UPF1, and RNA splicing factors) suggesting a role in the regulation of gene expression in response to DNA damage. Indeed, BCL11b overexpression was shown to repress DNA damages events (Cherrier et al., 2009). Then, we highlighted the potential role of BCL11b in autoregulation of some specific gene translation such as tubulin.

**Figure 7.**
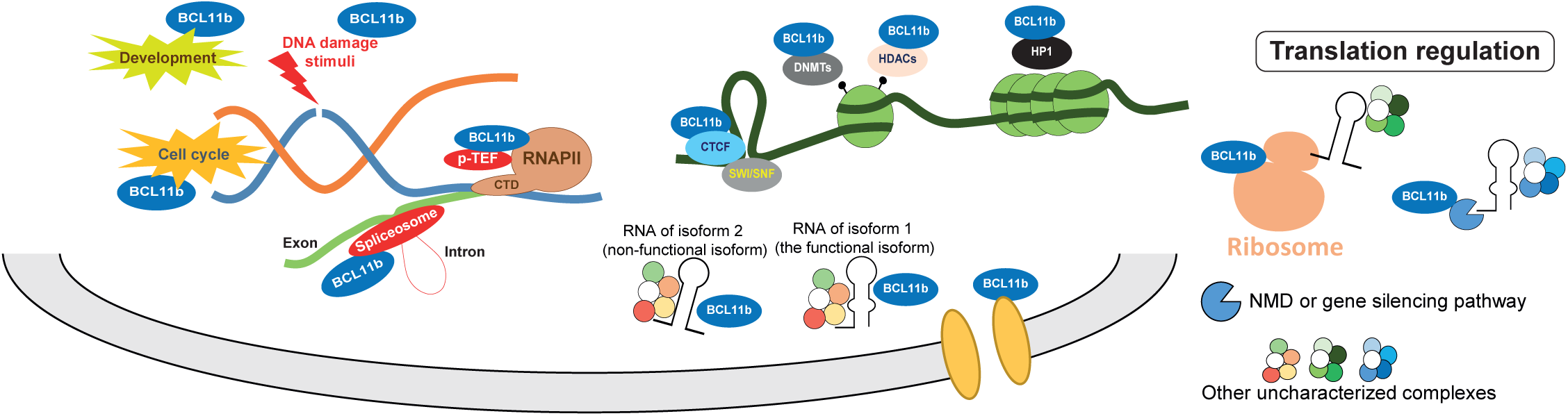
The graphical abstract shows the major proteins pathways, complexes and based on GO terms obtained after IP-MS and CLIP-seq experiments.

We found that BCL11b preferentially binds to RNA isoforms coding for proteins, in addition to BCL11b is associated with isoforms that do not code for proteins (INCPs), such as NMD and IR. Generally, translation is regulated by selecting the desired mRNA for being translated, whereas, the RNA decay pathways, such as Drosha, UPF1 (NMD pathway), and Ago (RNA interference) are able to decay or silence the undesirable RNA isoforms (Tuck et al., 2020; Wang and Aifantis, 2020). Our results suggest that BCL11b may contribute to the selection of RNA isoforms. Selecting the isoform (so-called isoform switching or selection) is a crucial process through which cells select a specific RNA isoform during the development or immune response. Some isoforms might be needed at specific steps during development, cell cycle or response to stimuli. Therefore, malfunction of RNA splicing or processing has a great impact on cell proliferation and cancer prognosis (Wang and Aifantis, 2020).

In conclusion, our analysis of BCL11b-protein and BCL11b-RNA interactions provide the first global overview of BCL11b interactants. The results suggest that BCL11b could bind to RNA processing proteins to regulate translation and/or select the desirable RNA isoform. One can speculate that depletion of BCL11b could lead to isoform switching and translation of wrong isoform of the protein (misfolded or short) but this needs to be formally demonstrated. Although the exact role of BCL11b in diseases deserve to be studied by future research, our results offer potential genome-wide RNA and proteins datasets, which will help future efforts to understand the molecular basis of cancer and neurodegenerative diseases.

## Human or animal experiments

no human or animal work have been performed

## Author contribution

HS conceived and designed bioinformatics analysis; HS, MDR, AAA and MK conceived and performed experiments; CW, TL, and FD provided scientific and logistic supports; CVL, CS and OR conceived the experiments, supervised the work and supported with resources. HS drafted the manuscript, all authors agreed for the final version of the manuscript.

Data deposited on SRA database with ID PRJNA661202

## Supporting information

Supplemental Informations

