## Supplemental Informations for "BCL11b interacts with RNA and proteins involved in RNA processing and developmental diseases"

### Supplementary files

**Supplementary file 1** (this file). This file, which contains supplementary material and methods, four supplementary tables (tables S1-S4), and supplementary figures (from figures S1-S10)

**External Supplementary Datasets.** In Excel sheet format (from dataset S1-S6)

**Supplementary Dataset 1**. Number of CLIP-seq reads mapped to human genome, and list of genes with significant BCL11b CLIP-seq reads in HEK293 cells and microglial cells.

**Supplementary Dataset 2**. GO terms of the significant CLIP-seq genes, KEGG pathways, gene set enrichment analysis (GSEA) and BCL11b-associated ribonucleoprotein complexes (spliceosome proteins and ncRNAs that bound to BCL11b).

**Supplementary Dataset 3**. Regions (10kb bins clusters and dCLIP tool) with highest number of CLIP RNA reads, and the associated GO terms and disease Ontology.

**Supplementary Dataset 4**. Number of CLIP-seq reads mapped to the transcripts of the selected 23 genes, which is used to construct Figure 2.

**Supplementary Dataset 5**. List BCL11b-protein interactants as deciphered by IP-MS, the complexes, GO terms, GSEA and KEGG pathways.

**Supplementary Dataset 6**. List of genes, which are shared between CLIP-seq with IP-MS, i.e. BCL11b binds to both protein products and mRNA transcripts of same gene, and GO.

### SUPPLEMENTARY TABLES

**Table S1**. The type of BCL11b-bound RNAs. Most of RNA transcripts are proteins coding.

| Gene type from Ensembl database | # of genes | % |
|---|---|---|
| protein_coding | 730 | 96.05% |
| processed_transcript | 3 | 0.39% |
| antisense | 3 | 0.39% |
| snRNA | 3 | 0.39% |
| rRNA | 2 | 0.26% |
| snoRNA | 6 | 0.79% |
| misc_RNA | 2 | 0.26% |
| pseudogene | 3 | 0.39% |
| Mt_rRNA | 1 | 0.13% |
| sense_intronic | 1 | 0.13% |
| miRNA | 1 | 0.13% |
| lincRNA | 5 | 0.66% |
| **Total** | **760** | |

**Table S2**. The involvement of BCL11b-bound RNAs and proteins in complexes. DE means differentially expressed in SMA or ALS diseases. For list of spliceosome proteins bound to BCL11b, see supplementary dataset 9. FUS-SMN1 complex, means the protein can bind to FUS and SMN1 to form large complex, see list in table S4.

<table>
<tr><th rowspan="2"></th><th rowspan="2">Number of proteins in complex</th><th colspan="3">Bound to BCL11b</th></tr>
<tr><th>Number of CLIP-seq genes and percent of BCL11b CLIP gene (%)</th><th>Number of proteins</th><th>% of complex</th></tr>
<tr><td>DE in SMA</td><td>-</td><td>316 (42%)</td><td>-</td><td>-</td></tr>
<tr><td>DE in ALS</td><td>-</td><td>47 (6.2%)</td><td>-</td><td>-</td></tr>
<tr><td>FUS-SMN1 complex</td><td>158</td><td></td><td>46</td><td>29%</td></tr>
<tr><td>Spliceosome</td><td>136</td><td rowspan="8">RNU5A-1 ncRNA</td><td>26</td><td>19%</td></tr>
<tr><td>Minor spliceosome</td><td>38</td><td>8</td><td>21%</td></tr>
<tr><td>Major spliceosome</td><td>244</td><td>27</td><td>11%</td></tr>
<tr><td>Spliceosomal A complex</td><td>19</td><td>9</td><td>47%</td></tr>
<tr><td>Spliceosomal C complex</td><td>53</td><td>10</td><td>19%</td></tr>
<tr><td>Spliceosomal P complex</td><td>49</td><td>11</td><td>22%</td></tr>
<tr><td>U5 small nucleolar ribonucleoprotein</td><td>9</td><td>5</td><td>56%</td></tr>
</table>

**Table S3**. Some BCL11b-interacting proteins were found to form complexes with FUS-SMN1 (158 protein), 46 proteins (29%) of this complex interact with BCL11b.

| UniProtKB ID | Gene name | UniProtKB ID | Gene name | UniProtKB ID | Gene name |
|---|---|---|---|---|---|
| O43823 | AKAP8 | P07910 | HNRNPC | Q99590 | SCAF11 |
| Q14444 | CAPRIN1 | Q14103 | HNRNPD | Q13435 | SF3B2 |
| O60563 | CCNT1 | O43390 | HNRNPR | P51532 | SMARCA4 |
| Q99459 | CDC5L | Q6PKG0 | LARP1 | Q92922 | SMARCC1 |
| P50750 | CDK9 | P46013 | MKI67 | Q16637 | SMN1 |
| Q14839 | CHD4 | P06748 | NPM1 | O75643 | SNRNP200 |
| Q92841 | DDX17 | P11940 | PABPC1 | P08621 | SNRNP70 |
| Q9UHI6 | DDX20 | Q86W92 | PPFIBP1 | Q07955 | SRSF1 |
| P17844 | DDX5 | Q6P2Q9 | PRPF8 | P84103 | SRSF3 |
| O43143 | DHX15 | P48634 | PRRC2A | O60506 | SYNCRIP |
| Q08211 | DHX9 | Q5JSZ5 | PRRC2B | P46937 | YAP1 |
| Q9C005 | DPY30 | Q9Y520 | PRRC2C | Q86VM9 | ZC3H18 |
| P35637 | FUS | Q14498 | RBM39 | Q6NZY4 | ZCCHC8 |
| P09651 | HNRNPA1 | Q9Y580 | RBM7 | Q96KR1 | ZFR |
| P51991 | HNRNPA3 | Q9Y265 | RUVBL1 | | |
| Q99729 | HNRNPAB | Q9Y230 | RUVBL2 | | |

**Table S4**. PCR primers used in the study

| Amplicon | Forward primer | Reverse primer | Chr:from-to |
|---|---|---|---|
| FUS_amp-1 | AGCAGTGGTGGCTATGAACC | GGGCCACCAAATTTATTGAA | chr16:31196439-31199649 |
| FUS_amp-2 | GGACCAAGGATCACGTCATGAC | ACTCCCTTCCTTCTCTATCCCC | chr16:31199653-31199799 |
| SFPQ_amp-1 | CGATCATCCACTATTACAACAGC | TGAGGAGGCCTGGAGAGAAA | chr1:35656394-35657105 |
| SFPQ_amp-2 | GCCGAATGGGCTACATGGAT | TTCCTCTAGGACCCTGTCCA | chr1:35650134-35653596 |
| SRRM2_amp-1 | GGCAGCTCTTTTGATCCTCAGC | GTGAGCGAGAACTGCTAGACTC | chr16:2807873-2808502 |
| SRRM2_amp-2 | ACCGCTAAGAGAGGGCGATC | AGAGCGGGACCTGCCCC | chr16:2811994-2812287 |

**Table S5**. Raw PCR Ct values

**Raw Ct values**

**Total**

| | Repl # 1 | Repl # 2 | Repl # 3 | Repl # 4 | Repl # 5 | Repl # 6 | Repl # 7 |
|---|---|---|---|---|---|---|---|
| **FUS_amp-1** | 30.47 | 31.46 | 31.05 | 30.68 | 31.46 | 30.26 | 29.81 |
| **FUS_amp-2** | 24.06 | 23.94 | 24.53 | | | | |
| **SRRM2_amp-1** | 23.69 | 23.55 | 21.23 | 22.50 | | | |
| **SRRM2_amp-2** | 22.74 | 22.01 | 23.80 | | | | |
| **SFPQ_amp-1** | 20.38 | 19.81 | 24.76 | | | | |
| **SFPQ_amp-2** | 23.69 | 24.16 | 23.28 | | | | |

**Ctip2**

| | Repl # 1 | Repl # 2 | Repl # 3 | Repl # 4 | Repl # 5 | Repl # 6 | Repl # 7 |
|---|---|---|---|---|---|---|---|
| **FUS_amp-1** | 32.09 | 32.85 | 32.46 | 32.14 | 32.34 | 30.60 | 31.28 |
| **FUS_amp-2** | 28.18 | 28.25 | 28.73 | | | | |
| **SRRM2_amp-1** | 24.02 | 24.24 | 24.91 | 24.94 | | | |
| **SRRM2_amp-2** | 26.56 | 26.63 | 27.65 | | | | |
| **SFPQ_amp-1** | 25.70 | 26.03 | 28.20 | | | | |
| **SFPQ_amp-2** | 29.67 | 30.14 | 29.97 | | | | |

**Mock**

| | Repl # 1 | Repl # 2 | Repl # 3 | Repl # 4 | Repl # 5 | Repl # 6 | Repl # 7 |
|---|---|---|---|---|---|---|---|
| **FUS_amp-1** | 35.61 | 35.61 | 34.65 | 35.35 | 35.35 | 31.22 | 34.22 |
| **FUS_amp-2** | 28.30 | 28.50 | 29.32 | | | | |
| **SRRM2_amp-1** | 26.56 | 26.97 | 27.42 | 27.91 | | | |
| **SRRM2_amp-2** | 29.51 | 29.64 | 29.13 | | | | |
| **SFPQ_amp-1** | 27.93 | 28.01 | 29.15 | | | | |
| **SFPQ_amp-2** | 30.90 | 30.38 | 31.14 | | | | |

### SUPPLEMENTARY FIGURES

**Figure S1**. BCL11b CLIP-seq tags (reads) of mapped to EWSR1, DNMT1, and SRRM2 genes. The results obtained from over-expression of BCL11b in HEK cells (a) and its control (b), as well as, endogenous BCL11b in microglial cells (c) and its control (d). In the genome browser screenshot below, additional reads represent other replicate and control and the reads alignment are shown. The olive green bars refer to the NCBI RefSeq gene. The vertical black bars represent the GENCODE transcripts of the stated genes, the vertical blue bars transcripts refer to the protein non-coding transcripts that harbor significant CLIP reads and their coordinates correspond to intron of RefSeq gene, these intronic regions are highlighted with light rose colored boxes. The light blue horizontal boxes refer to introns without significant BCL11b CLIP reads. The X-axis represents the coordinates of the gene and the chromosome number. The Y-axis represents the number of CLIP reads in each condition. Additional figures of the CLIP reads aligned to each genes are also shown.

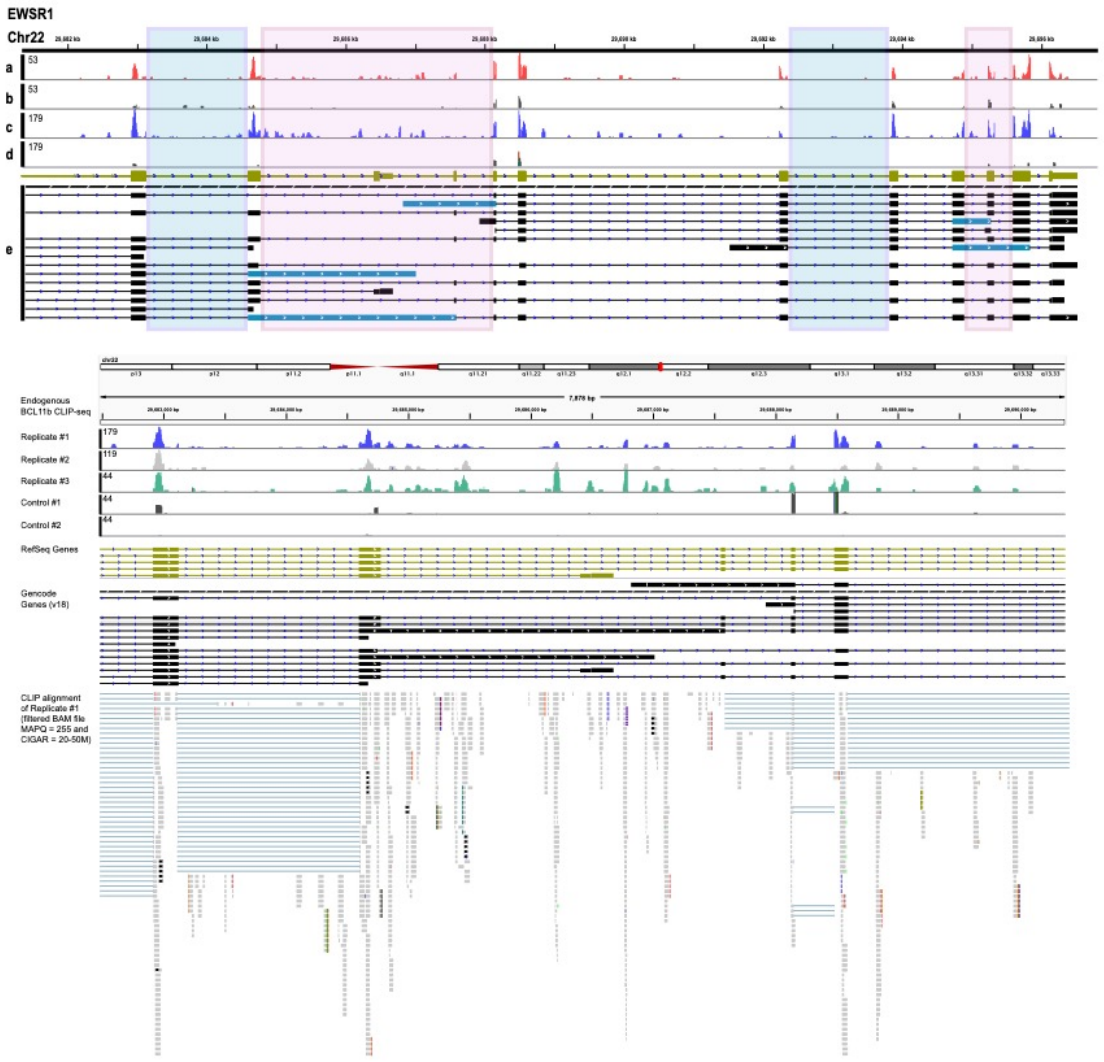

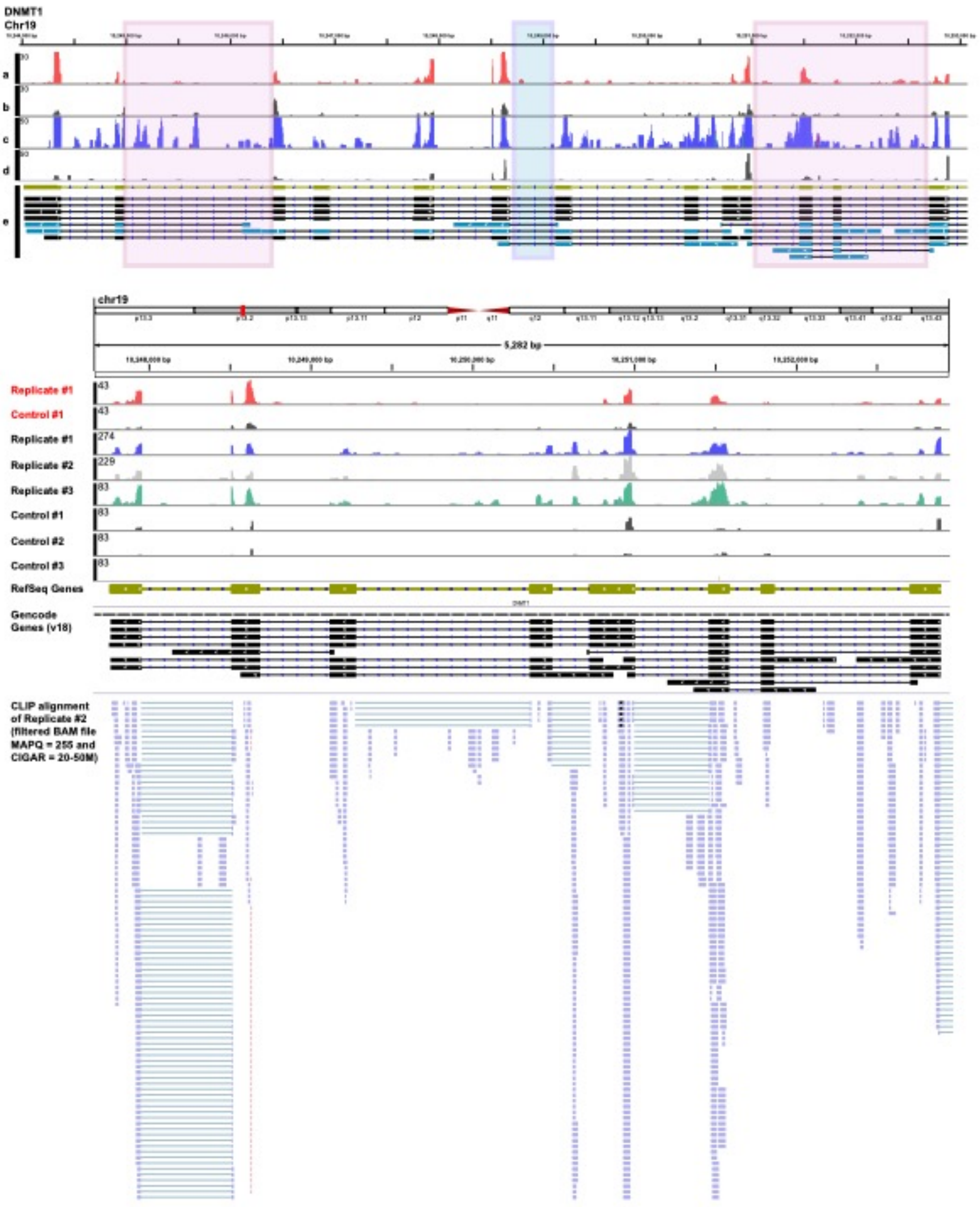
DNMT1
Chr19
a
b
c
d
e
chr19
5,282 bp
Replicate #1
Control #1
Replicate #1
Replicate #2
Replicate #3
Control #1
Control #2
Control #3
RefSeq Genes
Gencode Genes (v18)
CLIP alignment of Replicate #2 (filtered BAM file MAPQ = 255 and CIGAR = 20-50M)

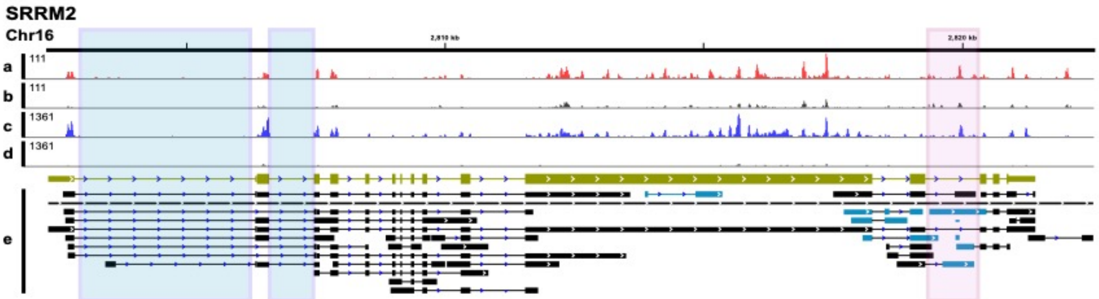
SRRM2
Chr16
2,810 kb
2,820 kb
a
111
b
111
c
1361
d
1361
e

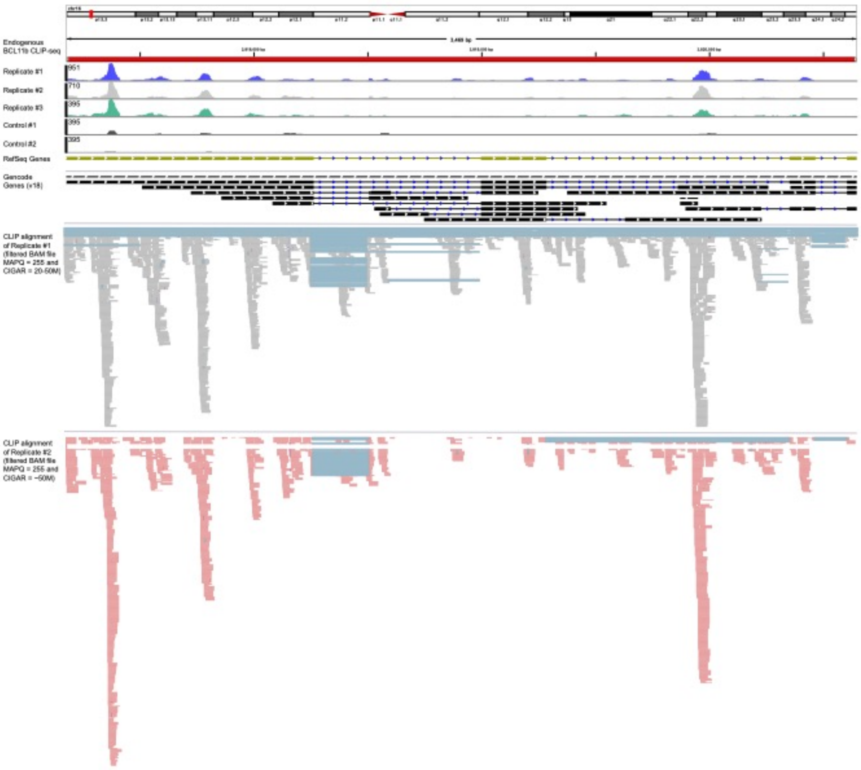
Endogenous BCL11b CLIP-seq
Replicate #1
951
Replicate #2
710
Replicate #3
395
Control #1
395
Control #2
395
RefSeq Genes
Gencode Genes (x18)
CLIP alignment of Replicate #1 (filtered BAM file MAPQ = 255 and CIGAR = 20-50M)
CLIP alignment of Replicate #2 (filtered BAM file MAPQ = 255 and CIGAR = ~50M)

**Figure S2**. The genes with top (high) CLIP-seq reads, i.e. high BCL11b-bound RNAs, which include ncRNA and protein-coding RNA. A) Top gene detected by CLIP-seq in HEK293 cells and (B) microglia. The volcano plot is constructed using R script. The values with "NA or undermined values" after or values with Zero CLIP-seq reads were omitted in both microglia and HEK cells, before run the R script.

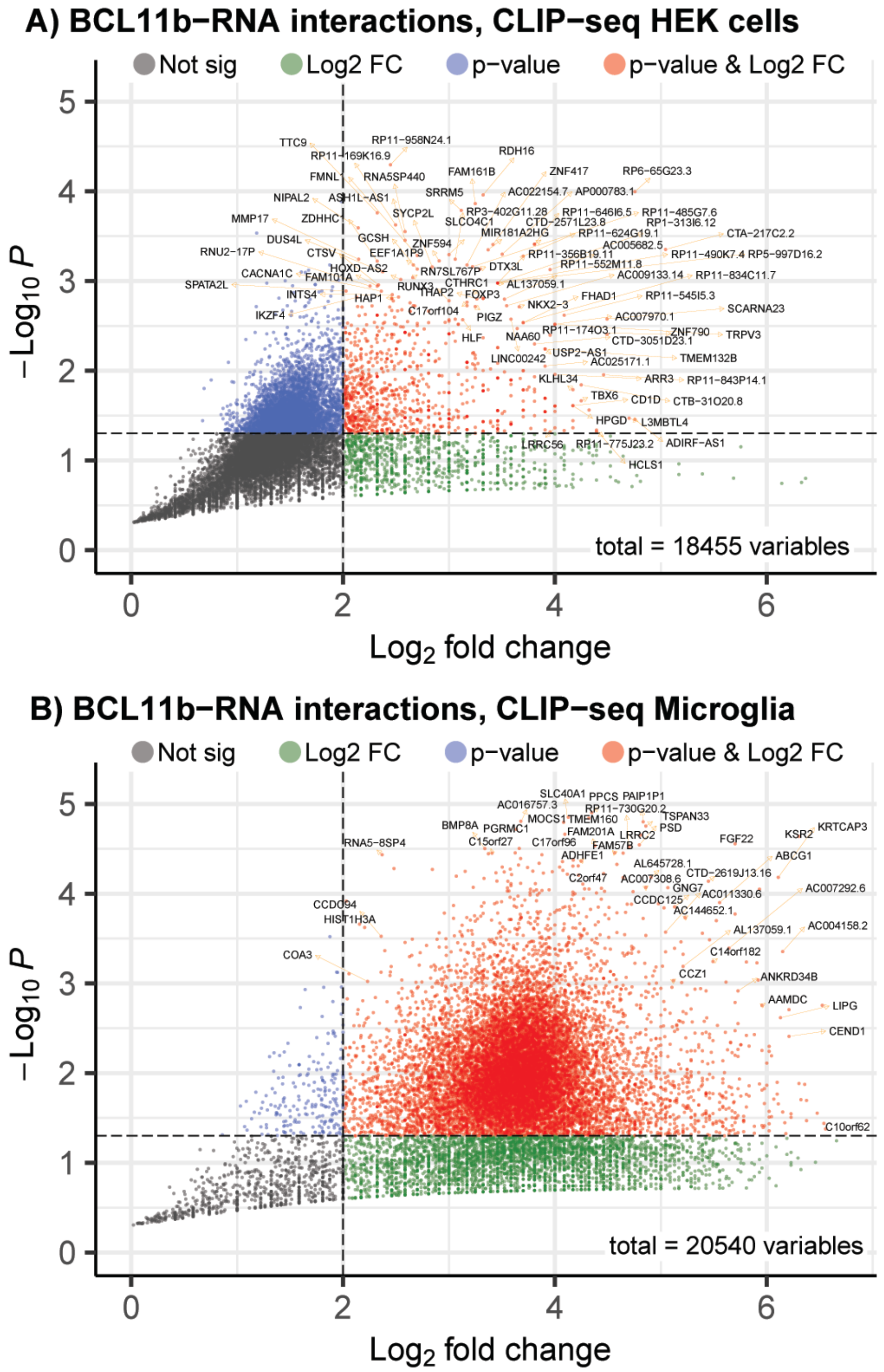

**Figure S3**. Functions of genes have significant CLIP-seq reads. (A) is GO terms of the genes that are significant in microglia and HEK cells. GSEA analysis of (B) endogenous CLIP-seq in microglia and (C) HEK-cells. (D) KEGG pathways of significant genes in HEK cells. Details can be found in Suppl. data S2

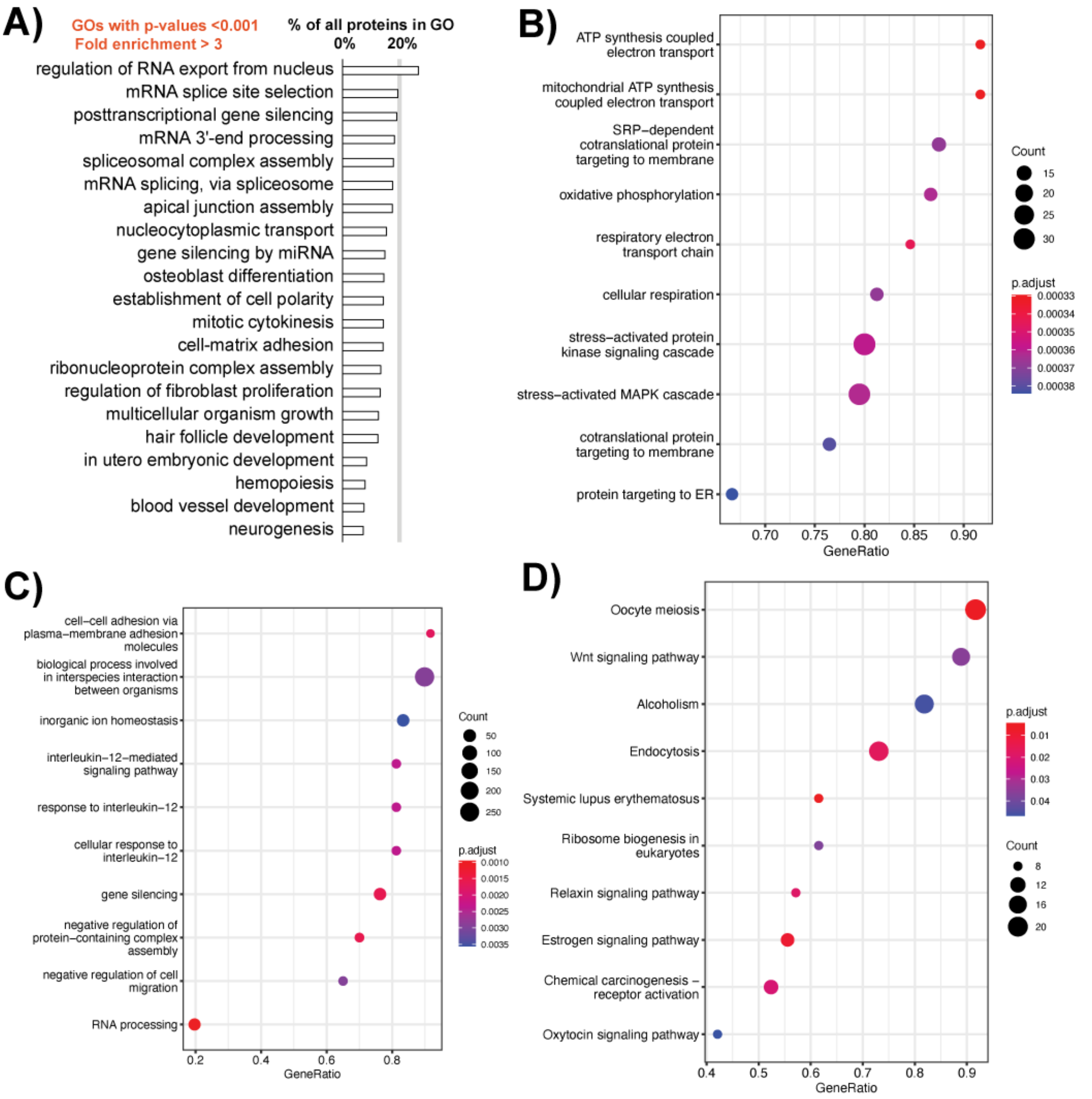

**Figure S4**. BCL11b binds to multiple RNA translated into protein in Wnt signaling patway CLIP-seq in microglia (in red boxes), and HEK cells (blue arrows). Comparing with previously published RNA-seq data, we found that the expression of most genes (and protein products) in the pathway are not changed as effect of BCL11b, example of RNA-seq in Hosokawa et al, 2018, doi: 10.1038/s41590-018-0238-4.

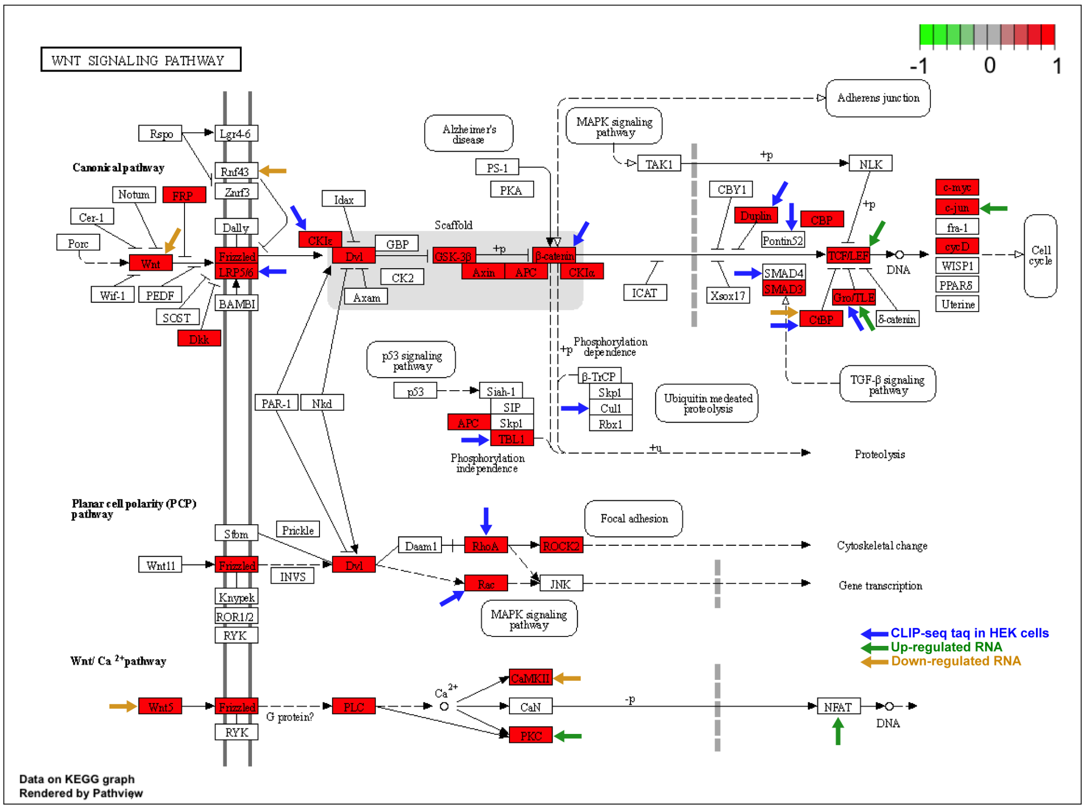

**Figure S5**. The locations of PCR amplicons used in RNA-IP-qPCR experiment, with respect to the gene coordinates, figure 4B and 4C in the main article, PCR primers in table S4.

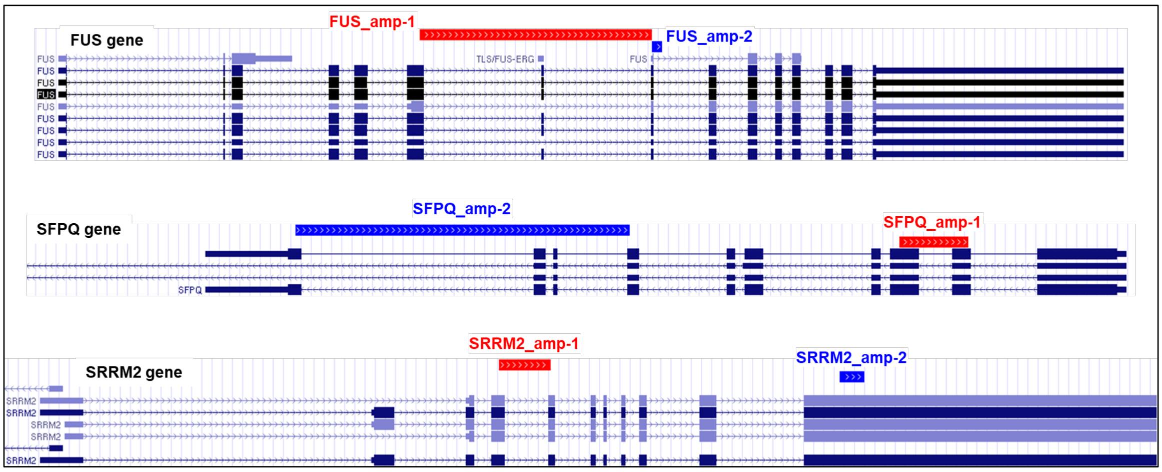


**Figure S6**. The q-rt-PCR results were urn into agarose gel. The gels show the PCR amplicons of figure 4B and 4C in the main article, PCR primers in table S4.

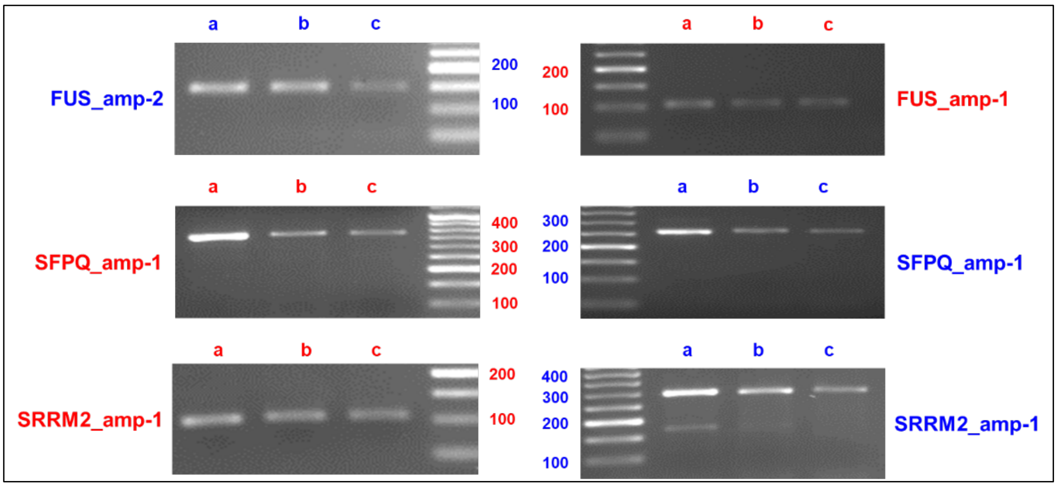

**Figure S7**. The location of CLIP clusters with respect to the TSS of genes. (A) 10kb bin clusters, and dCLIP results of (B) over-expressed BCL11b, and (C) endogenous BCL11b.

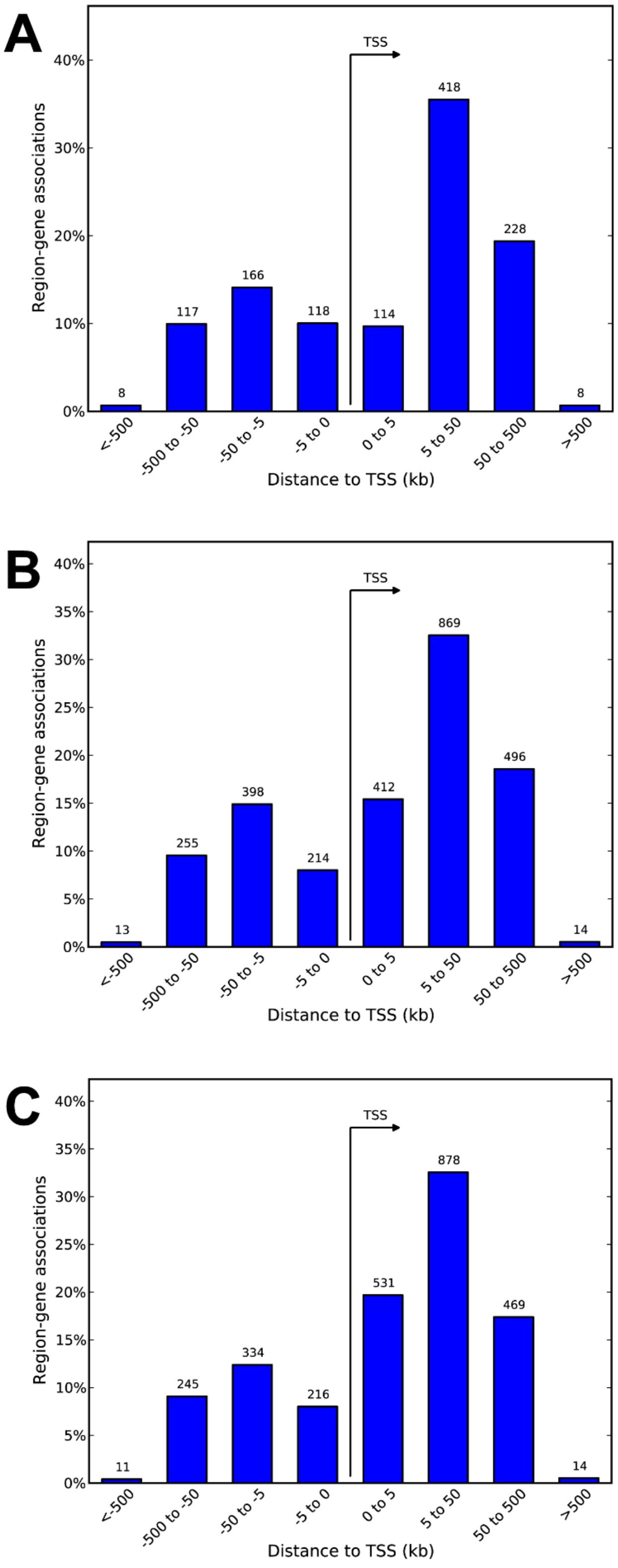

**Figure S8**. GO terms and disease annotation of BCL11b CLIP-seq reads. The CLIP reads clusters were counted per 10kb bins or using dCLIP tool (http://qbrc.swmed.edu/software/).

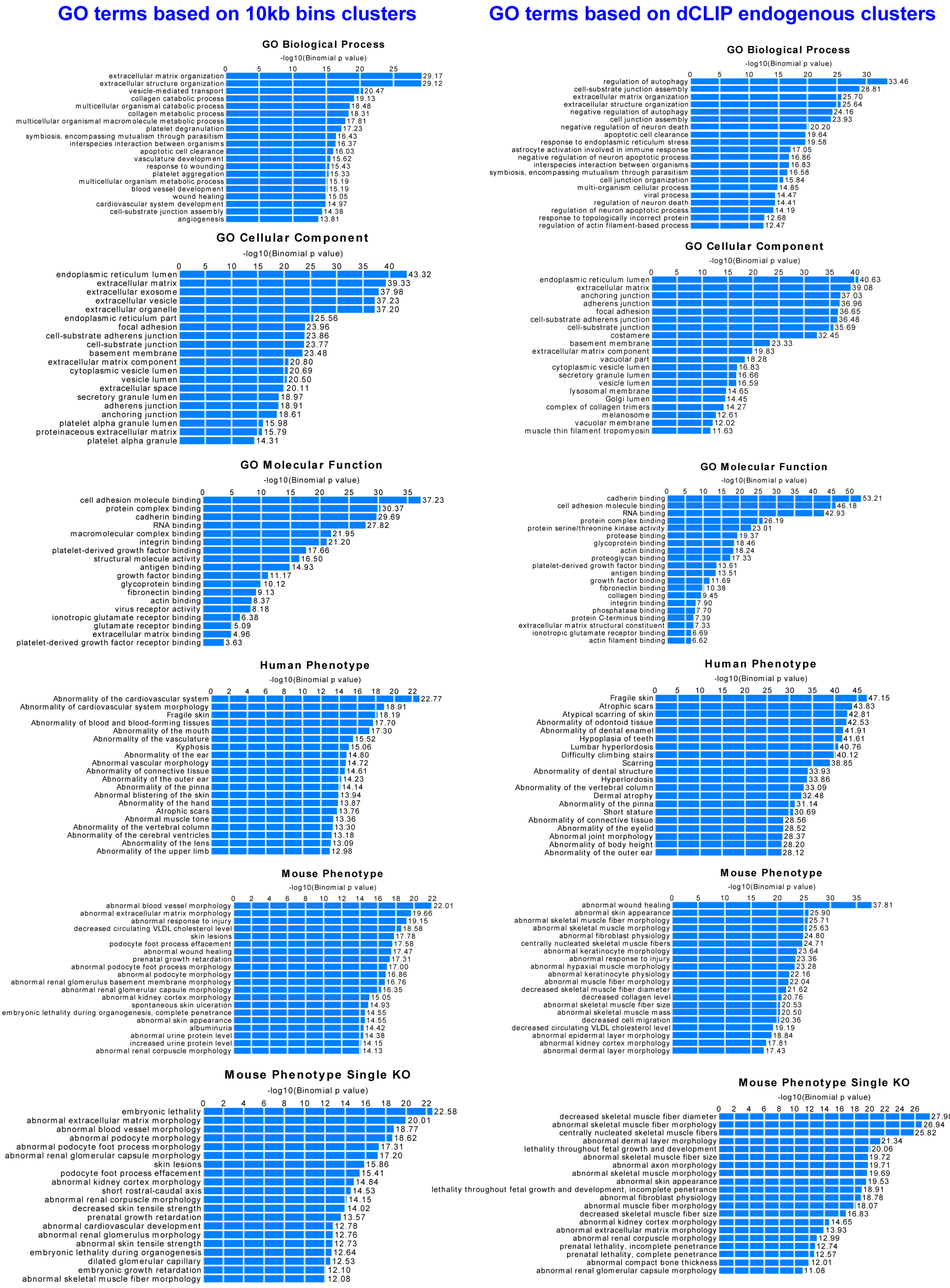

**Figure S9.** BCL11b-protein interactants (protein partner as identified by quantitative IP-MS. The top plot is the volcano plot of the top protein interactants that passed the cutoffs, whereas the boxplot below shows the median of 6 replicates (6 dots). The normalized log2FC shows proteins abundance (CTIP2/Control ratio) after CTIP2 overexpression.

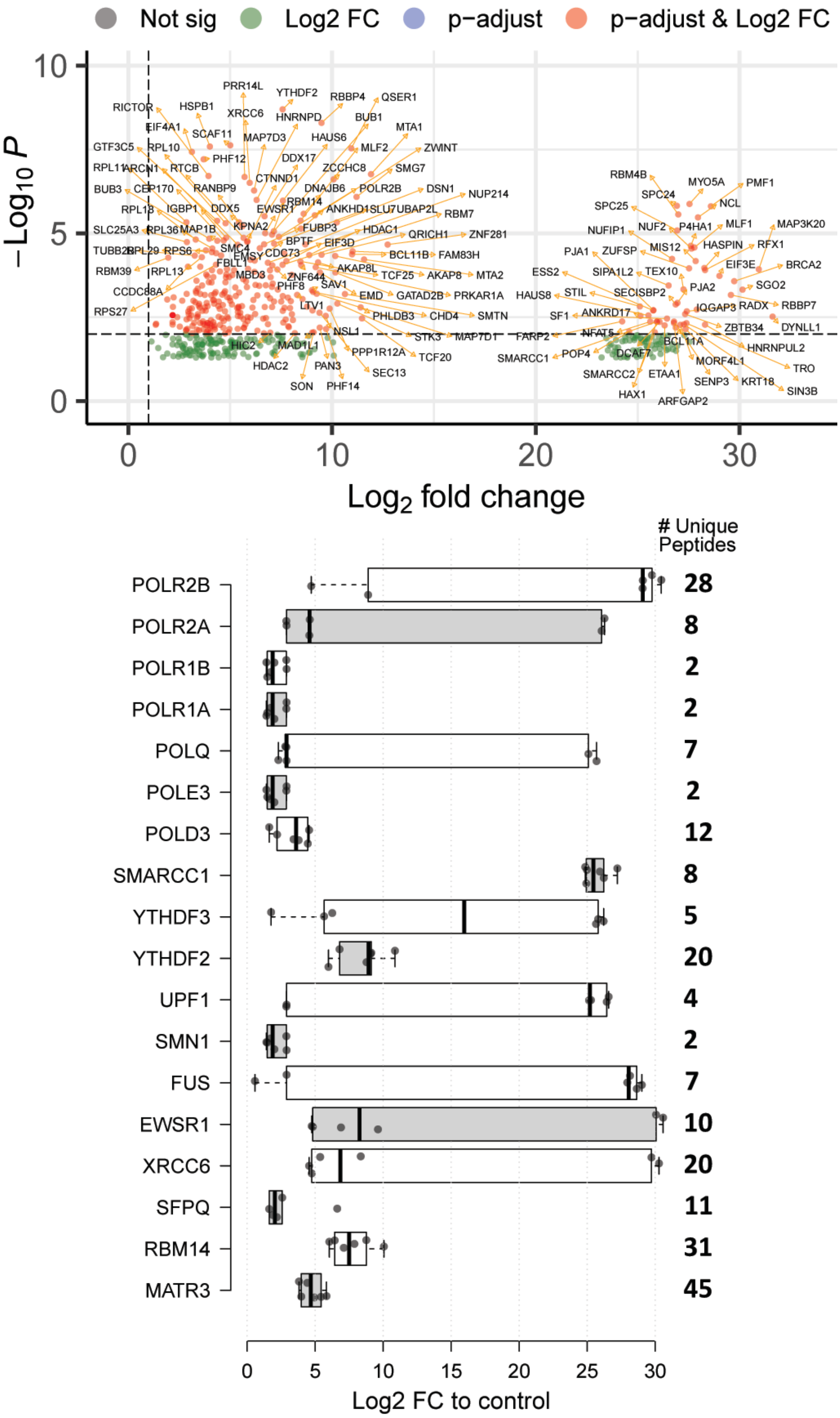

**Figure S10.** BCL11b binds to multiple complexes and pathways as deciphered by IP-MS results. A) In consistent with previous studies, using our IP-MS and CLIP-seq we can detect BCL11b interactions with proteins and ncRNA. B) GO terms of BCL11b-interacting proteins (BIPs), X-axis show percent of BIPs to the total number of proteins identified in this category. C) Gene set analysis (GSEA) of BCL11b-interacting proteins. D) Four representative GO terms with the list of proteins that are detected to bind to BCL11b. E) The complexes that constitute of BIPs, X-axis show percent of BIPs to the total number of proteins in the complex; the total number of proteins in the complex is written in dark blue outside the bars.

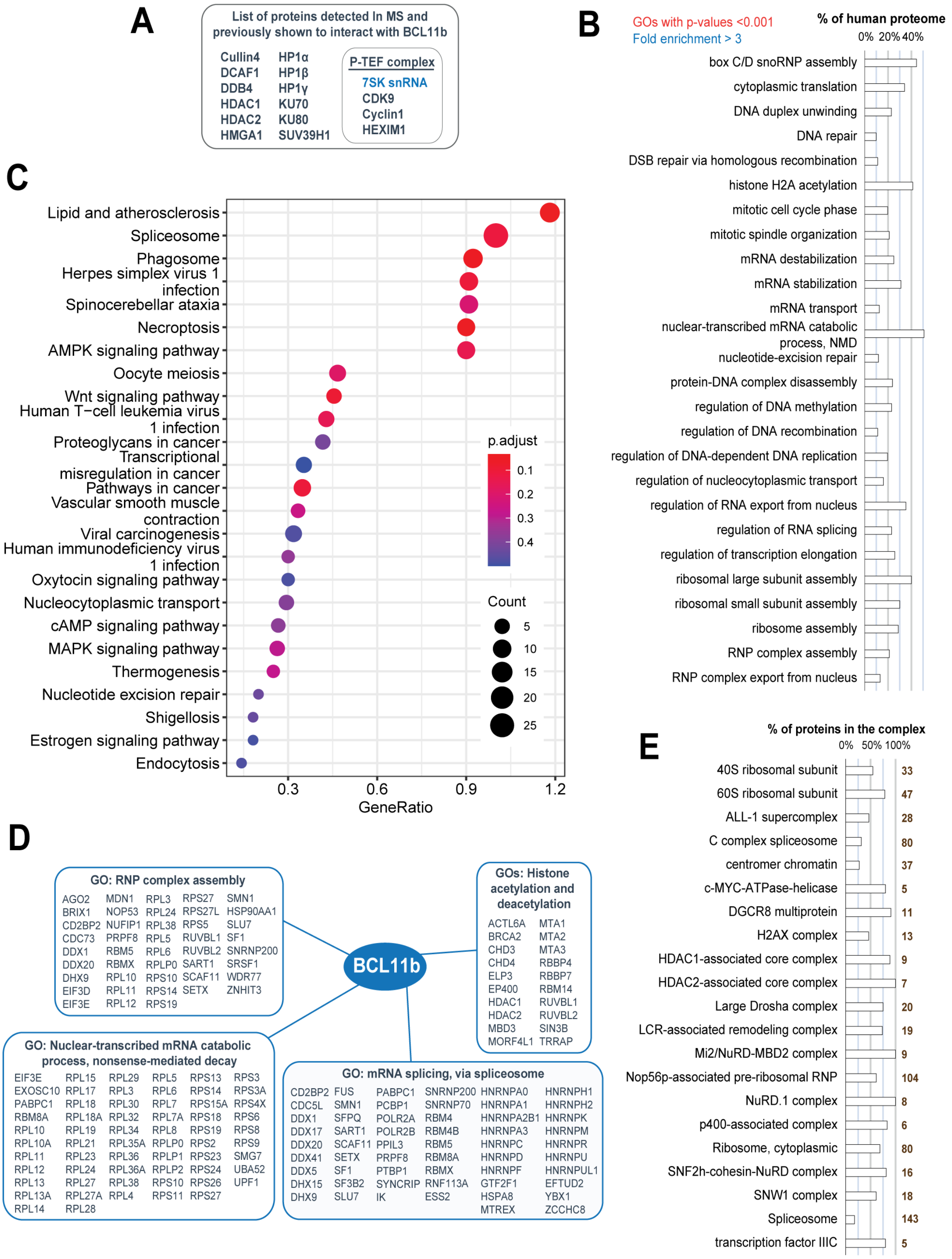

**Figure S11.** BCL11b binds to proteins (IP-MS) involved in cell cycle, KEGG ID: hsa04110.

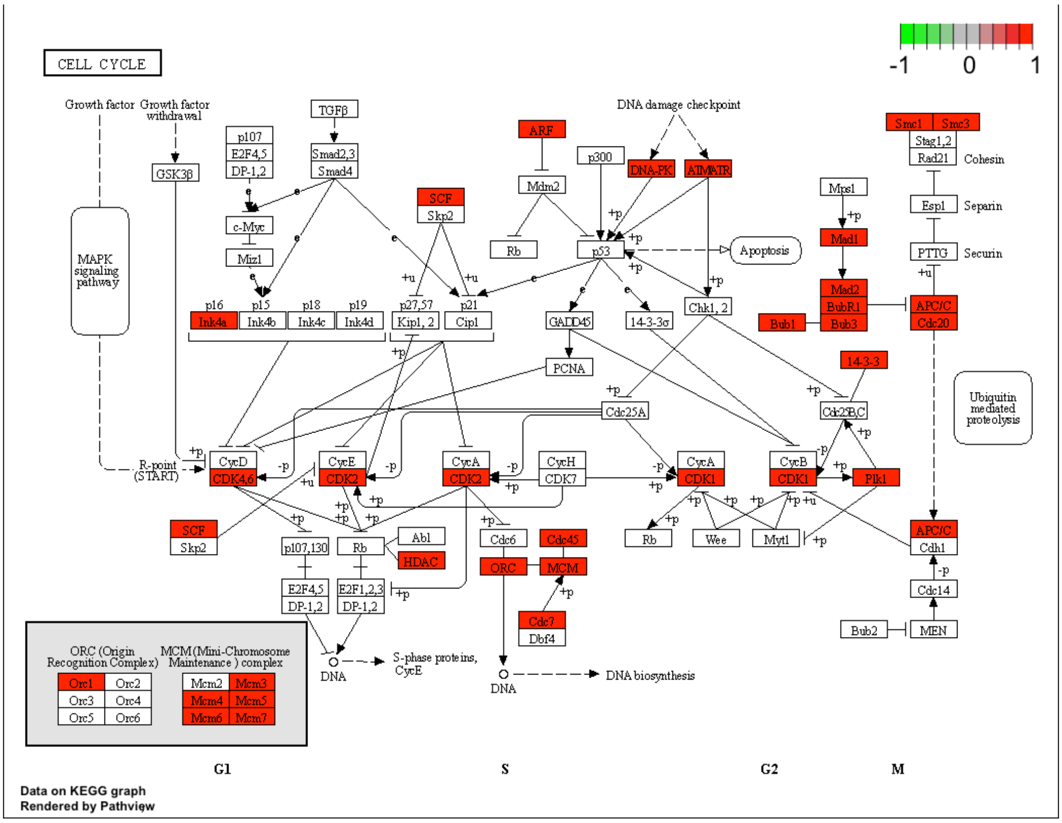

**Figure S12.** BCL11b binds to proteins (IP-MS) involved in RNA processing and metabolism pathways, KEGG IDs: (A) The ribosome biogenesis in eukaryotes “hsa03008”; (B) the mRNA surveillance pathway “hsa03015”; (C) RNA degradation “hsa03018”; and (D) the Spliceosome pathway “hsa03040”.

**A**

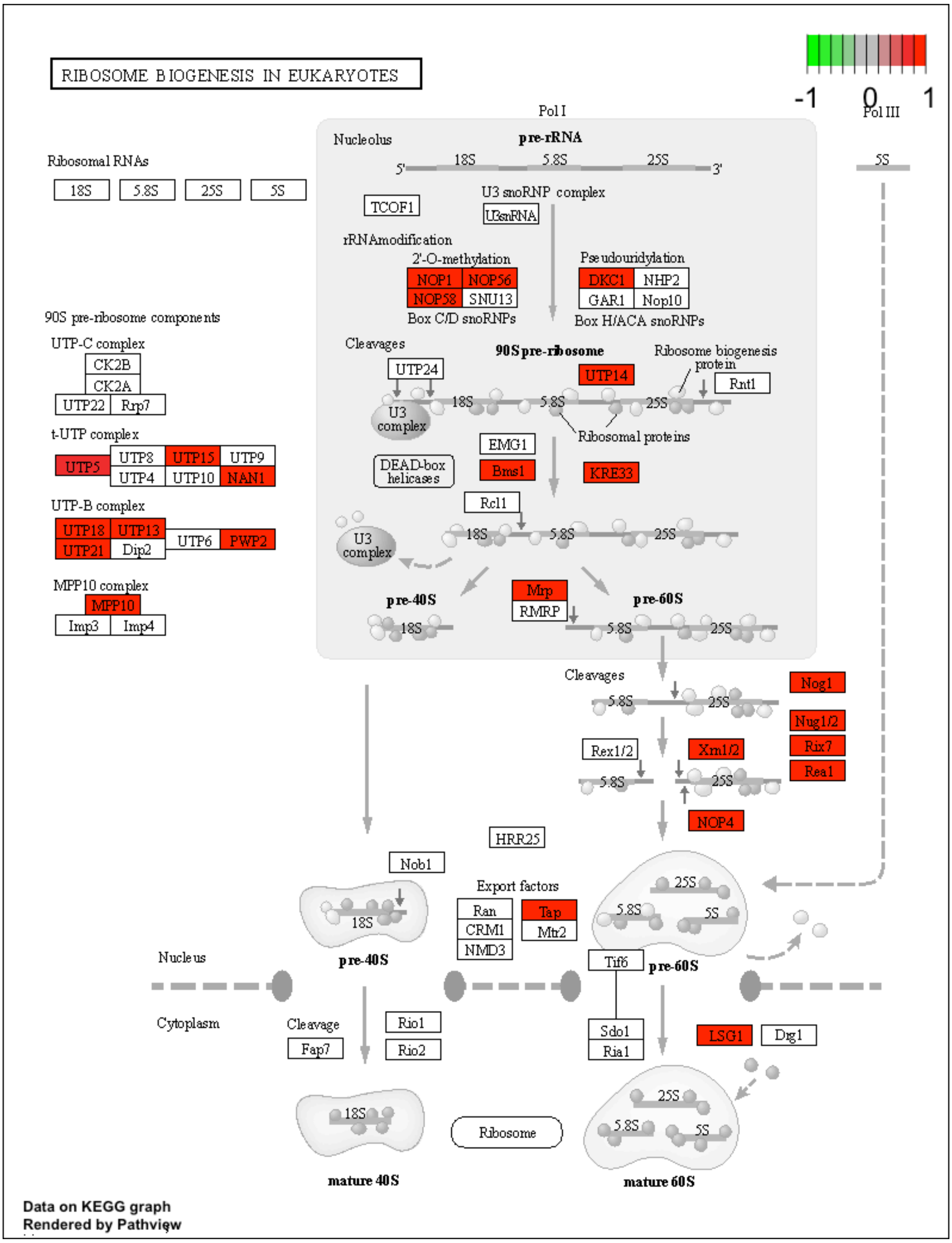

**B**

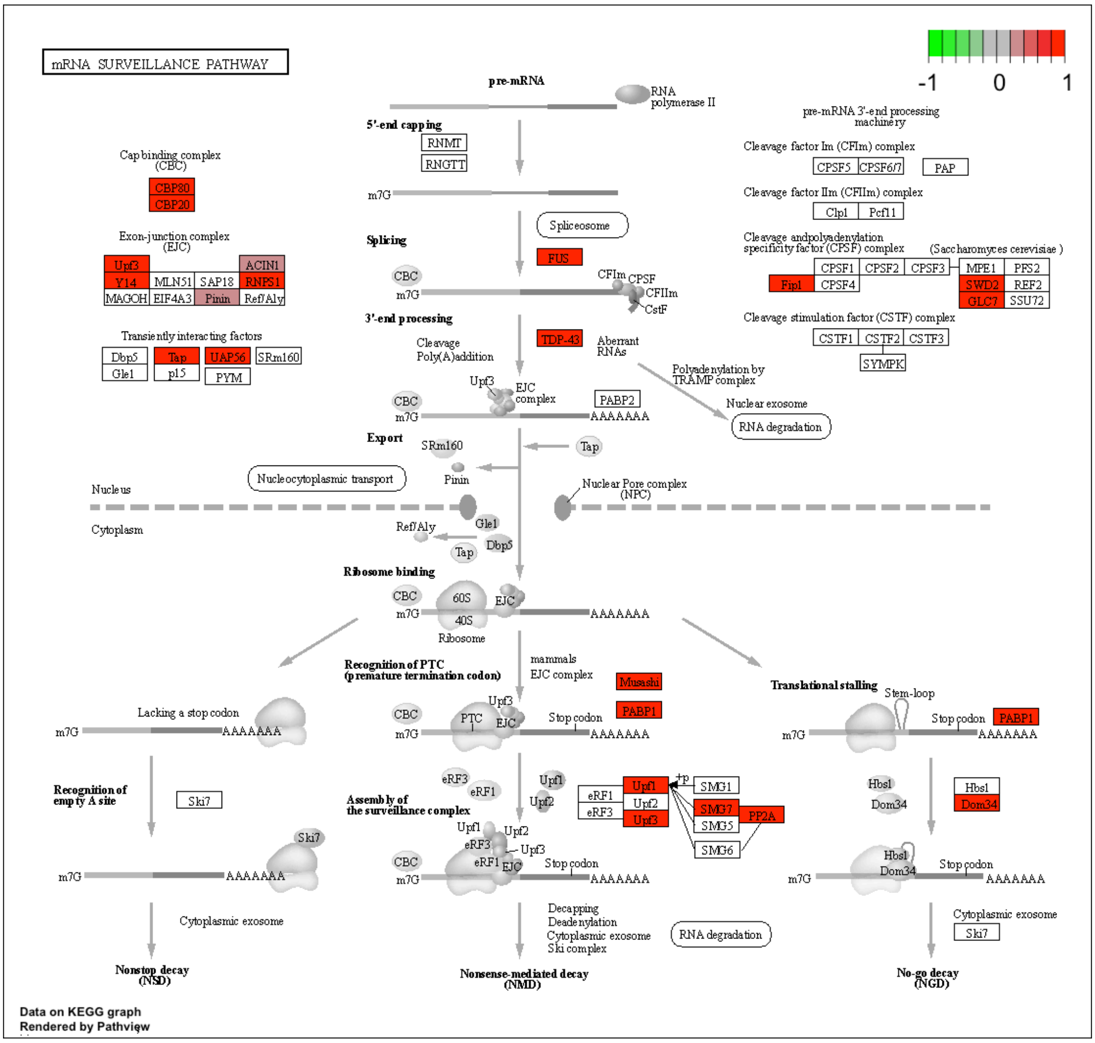
mRNA SURVEILLANCE PATHWAY
-1
0
1
pre-mRNA
RNA polymerase II
5'-end capping
RNMT
RNGTT
m7G
Splicing
Spliceosome
FUS
CBC
CFIm
CPSF
CFIIm
CstF
3'-end processing
TDP-43
Cleavage
Poly(A)addition
Aberrant RNAs
Upf3
EJC complex
PABP2
AAAAAAA
Export
SRm160
Tap
Nucleocytoplasmic transport
Pinin
Nuclear Pore complex (NPC)
Nucleus
Cytoplasm
Ref/Aly
Gle1
Dbp5
Ribosome binding
60S
40S
Ribosome
Cap binding complex (CBC)
CBP80
CBP20
Exon-junction complex (EJC)
Upf3
ACIN1
Y14
MLN51
SAP18
RNPS1
MAGOH
EIF4A3
Pinin
Ref/Aly
Transiently interacting factors
Dbp5
Tap
UAP56
SRm160
Gle1
p15
PYM
pre-mRNA 3'-end processing machinery
Cleavage factor Im (CFIm) complex
CPSF5
CPSF6/7
PAP
Cleavage factor IIm (CFIIm) complex
Clp1
Pcf11
Cleavage and polyadenylation specificity factor (CPSF) complex
(Saccharomyces cerevisiae)
CPSF1
CPSF2
CPSF3
MPE1
PFS2
Fip1
CPSF4
SWD2
REF2
GLC7
SSU72
Cleavage stimulation factor (CSTF) complex
CSTF1
CSTF2
CSTF3
SYMPK
Polyadenylation by TRAMP complex
Nuclear exosome
RNA degradation
Recognition of PTC (premature termination codon)
mammals
EJC complex
Musashi
PABP1
PTC
Stop codon
Lacking a stop codon
Recognition of empty A site
Ski7
eRF3
eRF1
Upf1
Upf2
Assembly of the surveillance complex
+p
SMG1
SMG7
PP2A
SMG5
SMG6
Decapping
Deadenylation
Cytoplasmic exosome
Ski complex
Translational stalling
Stem-loop
Hbs1
Dom34
Cytoplasmic exosome
Nonstop decay (NSD)
Nonsense-mediated decay (NMD)
No-go decay (NGD)
Data on KEGG graph
Rendered by Pathview

**C**

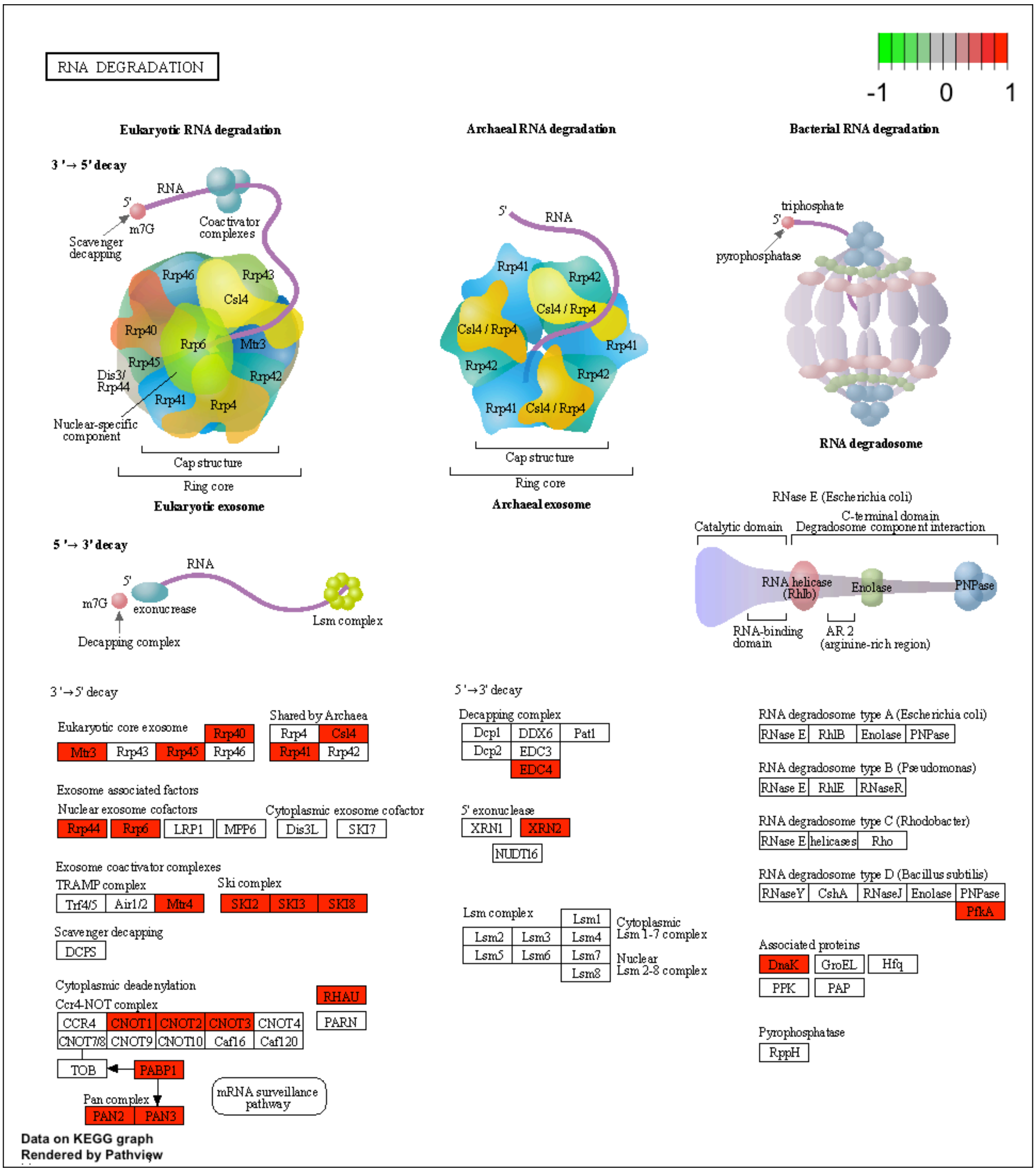
RNA DEGRADATION
-1 0 1
Eukaryotic RNA degradation
Archaeal RNA degradation
Bacterial RNA degradation
3'→5' decay
5'→3' decay
Eukaryotic exosome
Archaeal exosome
RNA degradosome
Cap structure
Ring core
RNase E (Escherichia coli)
Eukaryotic core exosome
Rrp40 Mtr3 Rrp43 Rrp45 Rrp46
Shared by Archaea
Rrp4 Csl4 Rrp41 Rrp42
Exosome associated factors
Nuclear exosome cofactors
Rrp44 Rrp6 LRP1 MPP6
Cytoplasmic exosome cofactor
Dis3L SKI7
Exosome coactivator complexes
TRAMP complex
Trf4/5 Air1/2 Mtr4
Ski complex
SKI2 SKI3 SKI8
Scavenger decapping
DCPS
Cytoplasmic deadenylation
Ccr4-NOT complex
CCR4 CNOT1 CNOT2 CNOT3 CNOT4
CNOT7/8 CNOT9 CNOT10 Caf16 Caf120
RHAU PARN
TOB PABP1
Pan complex
PAN2 PAN3
mRNA surveillance pathway
Decapping complex
Dcp1 DDX6 Patl
Dcp2 EDC3 EDC4
5' exonuclease
XRN1 XRN2 NUDT16
Lsm complex
Lsm1 Lsm2 Lsm3 Lsm4 Lsm5 Lsm6 Lsm7 Lsm8
Cytoplasmic Lsm 1-7 complex
Nuclear Lsm 2-8 complex
RNA degradosome type A (Escherichia coli)
RNase E RhlB Enolase PNPase
RNA degradosome type B (Pseudomonas)
RNase E RhlE RNaseR
RNA degradosome type C (Rhodobacter)
RNase E helicases Rho
RNA degradosome type D (Bacillus subtilis)
RNaseY CshA RNaseJ Enolase PNPase PfkA
Associated proteins
DnaK GroEL Hfq PPK PAP
Pyrophosphatase
RppH
Data on KEGG graph
Rendered by Pathview

**D**

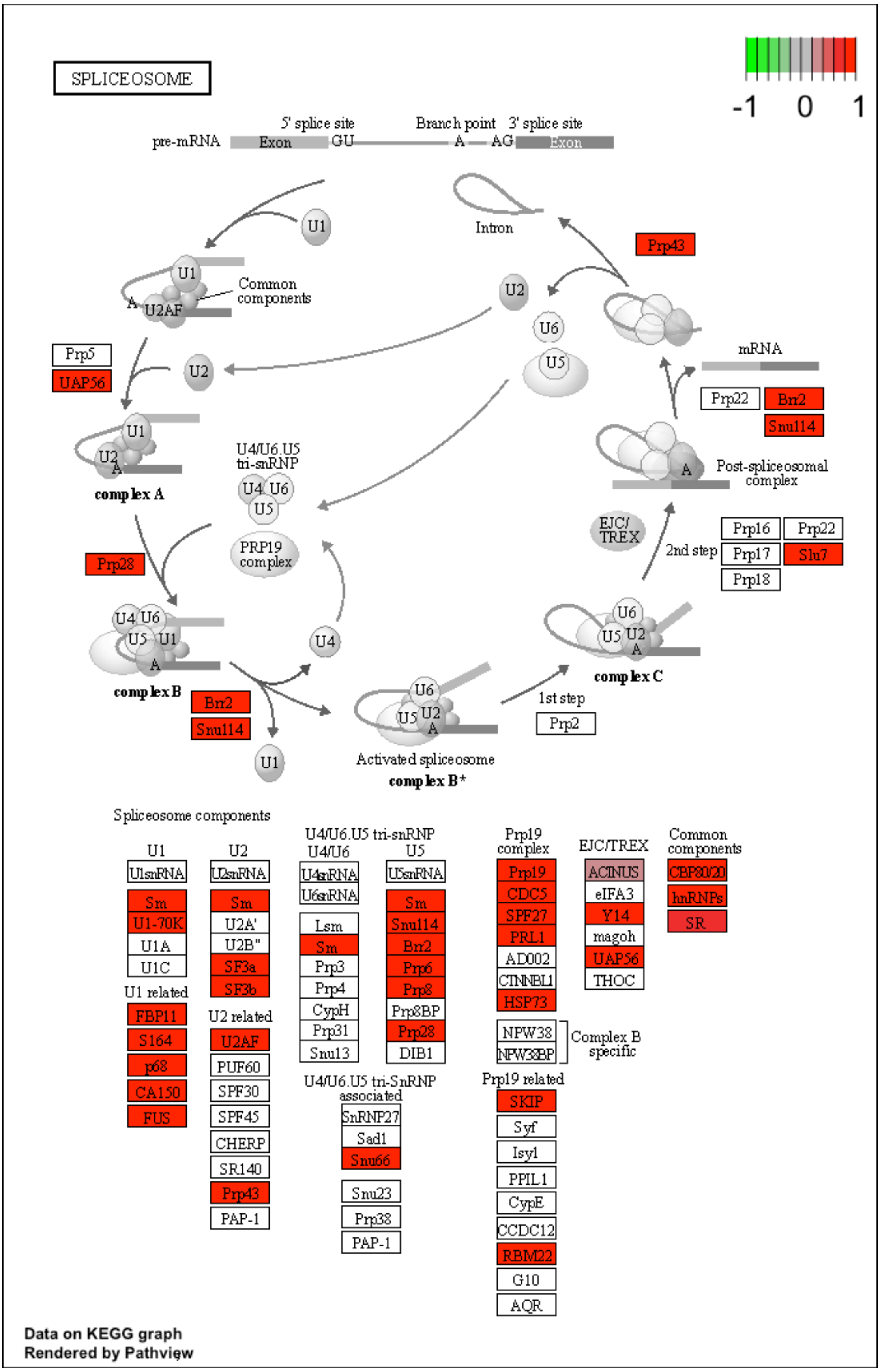
SPLICEOSOME
-1 0 1
5' splice site
Branch point
3' splice site
pre-mRNA
Exon
GU
A
AG
Exon
U1
Intron
Prp43
Common components
U2AF
U2
U6
U5
Prp5
UAP56
mRNA
Prp22
Brr2
Snu114
complex A
U4/U6.U5 tri-snRNP
U4 U6 U5
Post-spliceosomal complex
PRP19 complex
EJC/TREX
Prp16
Prp22
2nd step
Prp17
Slu7
Prp18
Prp28
U4
complex C
complex B
Brr2
Snu114
1st step
Prp2
Activated spliceosome
complex B*
Spliceosome components
U1
U1snRNA
Sm
U1-70K
U1A
U1C
U1 related
FBP11
S164
p68
CA150
FUS
U2
U2snRNA
Sm
U2A'
U2B"
SF3a
SF3b
U2 related
U2AF
PUF60
SPF30
SPF45
CHERP
SR140
Prp43
PAP-1
U4/U6.U5 tri-snRNP
U4/U6
U4snRNA
U6snRNA
Lsm
Sm
Prp3
Prp4
CypH
Prp31
Snu13
U5
U5snRNA
Sm
Snu114
Brr2
Prp6
Prp8
Prp8BP
Prp28
DIB1
U4/U6.U5 tri-SnRNP associated
SnRNP27
Sad1
Snu66
Snu23
Prp38
PAP-1
Prp19 complex
Prp19
CDC5
SPF27
PRL1
AD002
CTNNBL1
HSP73
NPW38
NPW38BP
Complex B specific
Prp19 related
SKIP
Syf
Isy1
PPIL1
CypE
CCDC12
RBM22
G10
AQR
EJC/TREX
ACINUS
eIFA3
Y14
magoh
UAP56
THOC
Common components
CBP80/20
hnRNPs
SR
Data on KEGG graph
Rendered by Pathview

**Figure S13.** BCL11b-binding RNA (microglial CLIP-seq data) encodes for proteins involved in cancer and leukemia pathways, KEGG IDs: (A) The Chronic myeloid leukemia pathway “hsa05220”; and (B) pathways in cancer “hsa05200”.

**A**

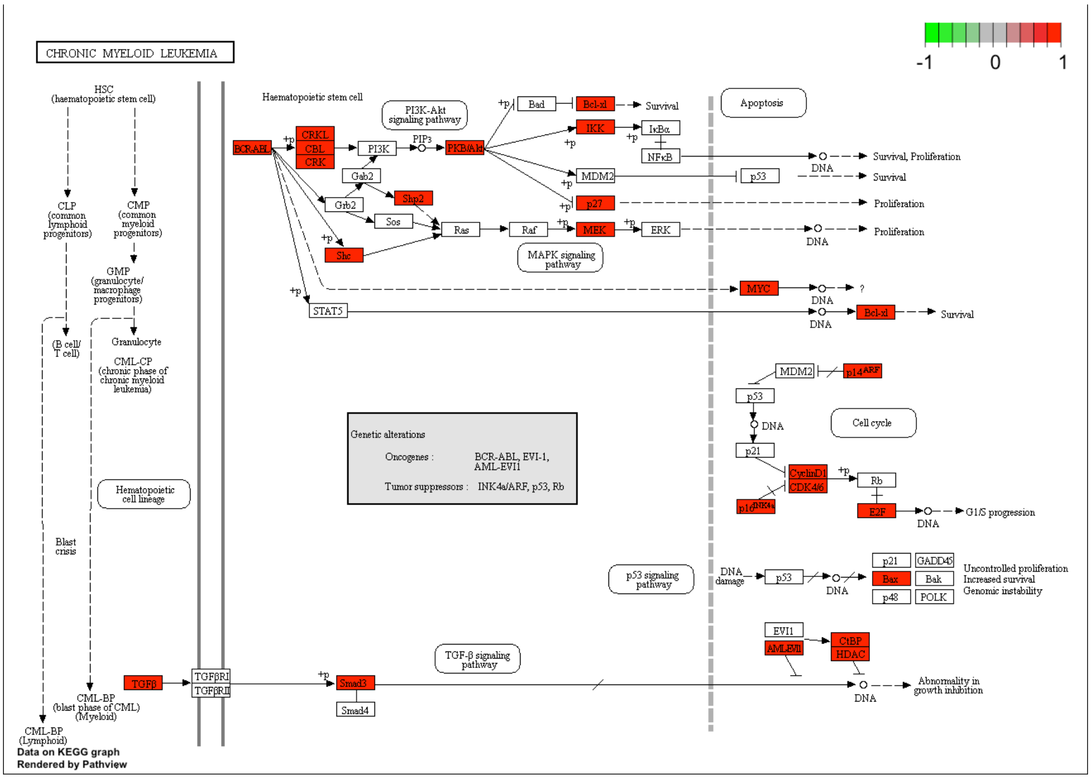

**B**

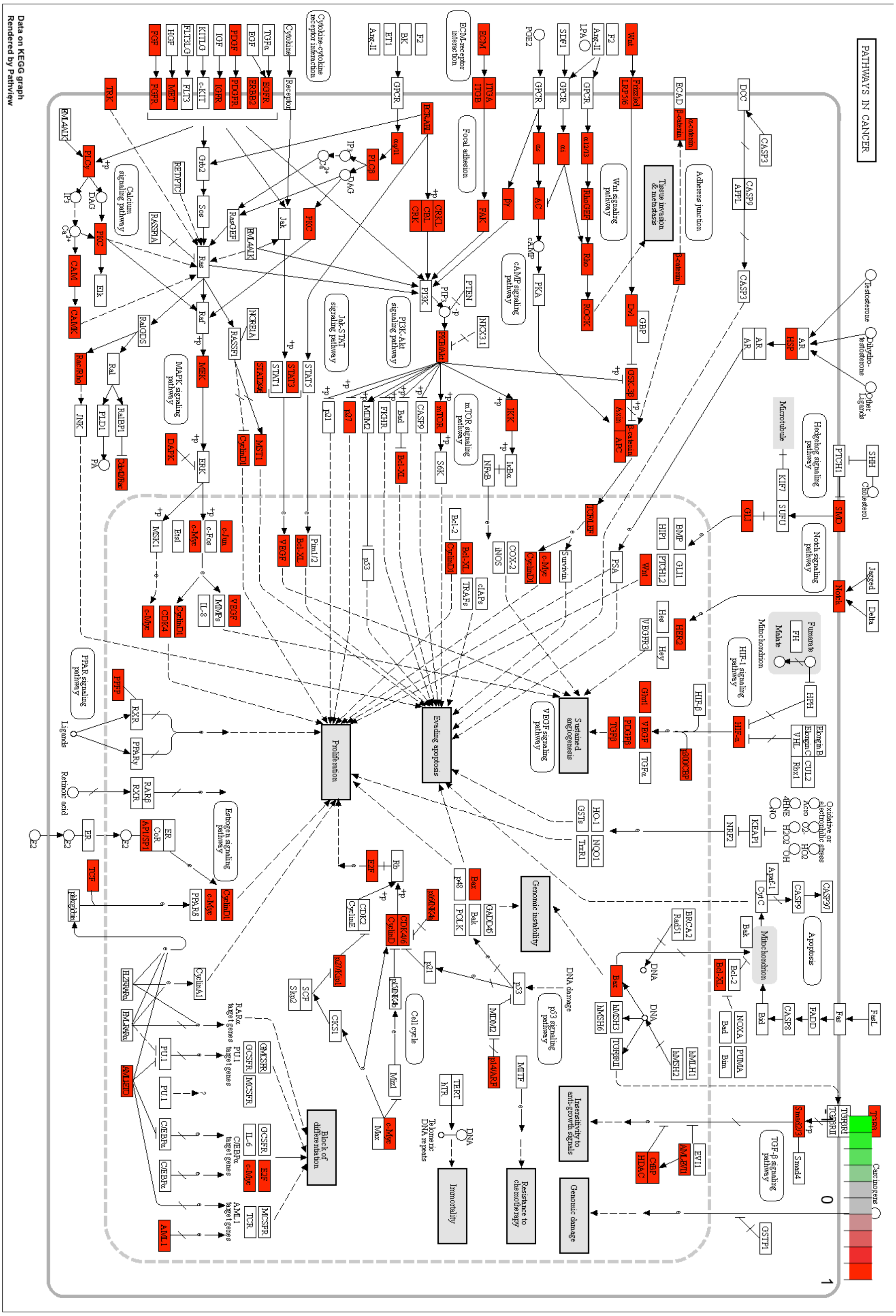

**Figure S14.** BCL11b-binding RNA (microglial CLIP-seq data) encodes for proteins involved in neurodegeneration, such as ALS and Huntington disease, KEGG IDs: (A) The amyotrophic lateral sclerosis pathway "hsa05014"; and the Huntington disease pathway "hsa05016".

**A**

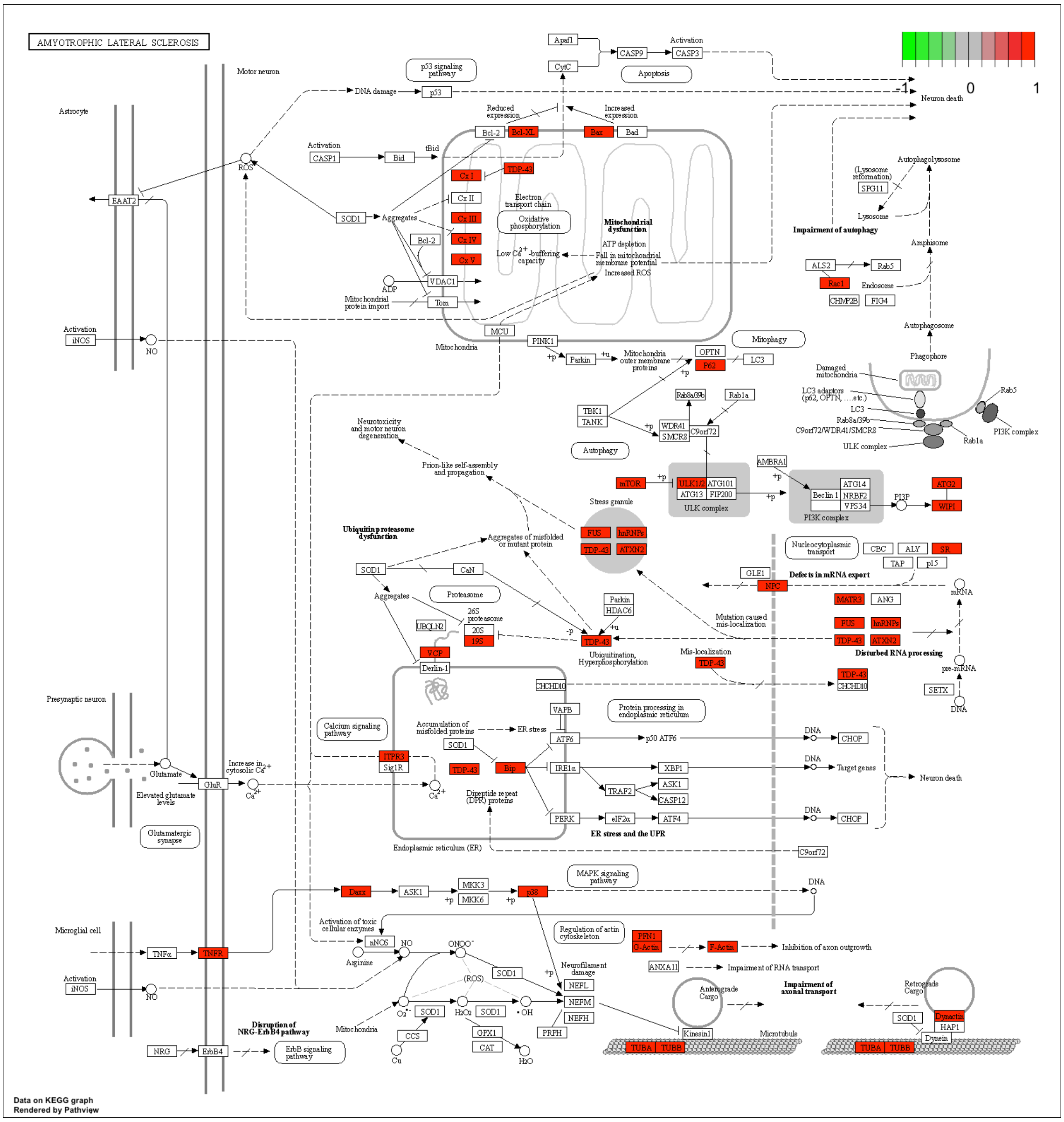

**B**

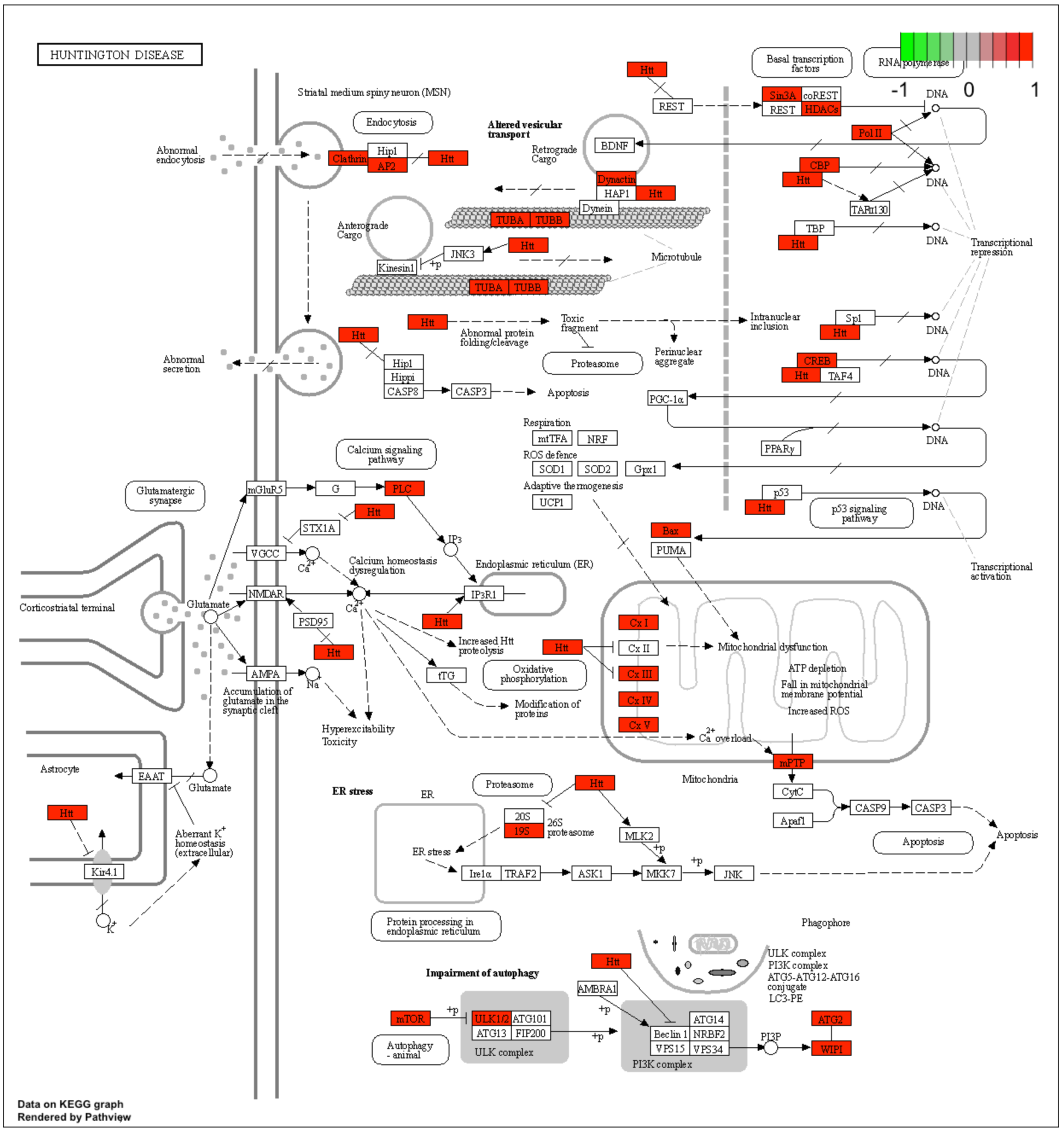
HUNTINGTON DISEASE
-1
0
1
Striatal medium spiny neuron (MSN)
Endocytosis
Abnormal endocytosis
Clathrin
Hip1
AP2
Htt
Altered vesicular transport
Retrograde Cargo
BDNF
Dynactin
HAP1
Dynein
TUBA
TUBB
Anterograde Cargo
JNK3
Kinesin1
Microtubule
Abnormal protein folding/cleavage
Toxic fragment
Proteasome
Perinuclear aggregate
Intranuclear inclusion
Abnormal secretion
Hippi
CASP8
CASP3
Apoptosis
Basal transcription factors
RNA polymerase
REST
Sin3A
coREST
HDACs
Pol II
CBP
TAFII130
TBP
Sp1
CREB
TAF4
PGC-1α
PPARγ
p53
p53 signaling pathway
DNA
Transcriptional repression
Transcriptional activation
Respiration
mtTFA
NRF
ROS defence
SOD1
SOD2
Gpx1
Adaptive thermogenesis
UCP1
Calcium signaling pathway
Glutamatergic synapse
mGluR5
G
PLC
IP3
STX1A
VGCC
Calcium homeostasis dysregulation
Endoplasmic reticulum (ER)
NMDAR
IP3R1
PSD95
Corticostriatal terminal
Glutamate
Increased Htt proteolysis
AMPA
tTG
Oxidative phosphorylation
Modification of proteins
Accumulation of glutamate in the synaptic cleft
Hyperexcitability
Toxicity
Bax
PUMA
Cx I
Cx II
Cx III
Cx IV
Cx V
Mitochondrial dysfunction
ATP depletion
Fall in mitochondrial membrane potential
Increased ROS
Ca overload
mPTP
Mitochondria
CytC
Apaf1
CASP9
Astrocyte
EAAT
Kir4.1
Aberrant K+ homeostasis (extracellular)
ER stress
ER
20S
19S
26S proteasome
MLK2
Ire1α
TRAF2
ASK1
MKK7
JNK
Protein processing in endoplasmic reticulum
Phagophore
ULK complex
PI3K complex
ATG5-ATG12-ATG16 conjugate
LC3-PE
Impairment of autophagy
AMBRA1
mTOR
ULK1/2
ATG101
ATG13
FIP200
Autophagy - animal
Beclin 1
ATG14
NRBF2
VPS15
VPS34
PI3P
ATG2
WIPI
Data on KEGG graph
Rendered by Pathview
